# fMRIflows: a consortium of fully automatic univariate and multivariate fMRI processing pipelines

**DOI:** 10.1101/2021.03.23.436650

**Authors:** Michael P. Notter, Peer Herholz, Sandra Da Costa, Omer F. Gulban, Ayse Ilkay Isik, Anna Gaglianese, Micah M. Murray

**Affiliations:** The Laboratory for Investigative Neurophysiology (The LINE), Department of Radiology, University Hospital Center and University of Lausanne, Switzerland; International Laboratory for Brain, Music and Sound Research, Université de Montréal & McGill University, Montréal, Canada; NeuroDataScience - ORIGAMI lab, McConnell Brain Imaging Centre, Montréal Neurological Institute, McGill University, Montréal, Canada; CIBM Center for Biomedical Imaging, Lausanne, Switzerland; Department of Cognitive Neuroscience, Maastricht University, The Netherlands; Brain Innovation B.V., Maastricht, The Netherlands; Department of Neuroscience, Max Planck Institute for Empirical Aesthetics, Frankfurt am Main, Germany; The Sense Innovation and Research Center, Lausanne and Sion, Switzerland

**Keywords:** Python, neuroimaging, data processing, pipeline, reproducible research

## Abstract

How functional MRI (fMRI) data are analyzed depends on the researcher and the toolbox used. It is not uncommon that the processing pipeline is rewritten for each new dataset. Consequently, code transparency, quality control and objective analysis pipelines are important for improving reproducibility in neuroimaging studies. Toolboxes, such as Nipype and fMRIPrep, have documented the need for and interest in automated pre-processing analysis pipelines. Recent developments in data-driven models combined with high-resolution neuroimaging datasets have strengthened the need not only for a standardized preprocessing workflow but also for a reliable and comparable statistical pipeline. Here, we introduce fMRIflows: a consortium of fully automatic neuroimaging pipelines for fMRI analysis, which performs standard preprocessing, as well as 1st- and 2nd-level univariate and multivariate analyses. In addition to the standardized pre-processing pipelines, fMRIflows provides flexible temporal and spatial filtering to account for datasets with increasingly high temporal resolution and to help appropriately prepare data for advanced machine learning analyses, improving signal decoding accuracy and reliability. This paper *first* describes fMRIflows’ structure and functionality, *then* explains its infrastructure and access, and *lastly* validates the toolbox by comparing it to other neuroimaging processing pipelines such as fMRIPrep, FSL and SPM. This validation was performed on three datasets with varying temporal sampling and acquisition parameters to prove its flexibility and robustness. fMRIflows is a fully automatic fMRI processing pipeline that uniquely offers univariate and multivariate single-subject and group analyses as well as pre-processing.

## 1 Introduction

Functional magnetic resonance imaging (fMRI) is a well-established neuroimaging method used to analyze activation patterns in order to understand brain function. A full fMRI analysis includes preprocessing of the data, followed by statistical analysis and inference of the results, usually separated into 1^st^-level analysis (the statistical analysis within subjects) and 2^nd^-level analysis (the group analysis between subjects). The goal of preprocessing is to identify and remove nuisance sources, measure confounds, apply temporal and spatial filters and to spatially realign and normalize images to make them spatially conform (Caballero-Gaudes and Reynolds, 2017). A good preprocessing pipeline should improve the signal-to-noise ratio (SNR) of the data, ensure the validity of inference and interpretability of results (Ashburner, 2009), reduce false positive and false negative errors in the statistical analysis and therefore improve the statistical power.

Even though the consequences of inappropriate preprocessing and statistical inference are well documented (Strother, 2006; Power et al., 2017b), most fMRI analysis pipelines are still established ad-hoc, subjectively customized by researchers to each new dataset (Carp, 2012). This usage can be explained by the circumstance that most researchers, by habit or lack of time, stick with the neuroimaging software at-hand or reuse and modify scripts and code snippets from colleagues and previous projects, and do not always adapt their processing pipelines to the newest standard in neuroimaging processing. Rehashing processing pipelines is associated with problems like persisting bugs in the code and delays in updating individual analysis steps to the most recent standards. This can lead to far-reaching consequences. Of course, the constant updating of pipelines to the newest standards and software also bears the risk of introducing new bugs and might lead to the pitfall of blindly trusting new untested procedures.

One solution to tackle this issue will require code transparency, good quality control and collective development of well-tested objective analysis pipelines (Gorgolewski et al., 2016). Recent years have brought some important reformations to the neuroimaging community that go in this direction.

*First*, the introduction of Nipype (Gorgolewski et al., 2011) made it easier for researchers to switch between different neuroimaging toolboxes, such as AFNI (Cox and Hyde, 1997), ANTs (Avants et al., 2011), FreeSurfer (Fischl, 2012), FSL (Jenkinson et al., 2012), and SPM (Friston et al., 2006). Nipype together with other software packages such as Nibabel (Brett et al., 2018) and Nilearn (Abraham et al., 2014) opened up the whole Python ecosystem to the neuroimaging community. Code can be shared between researchers via online services such as GitHub (https://github.com), and the whole neuroimaging software ecosystem can be run on any machine or server through the use of container software such as Docker (https://www.docker.com) or Singularity (https://www.sylabs.io). Combined with a continuous integration service such as CircleCI (https://circleci.com) or TravisCI (https://travis-ci.org), this allows the creation of easy-to-read, transparent, shareable and continuously tested open-source neuroimaging processing pipelines.

*Second*, the next major advancement in the neuroimaging field was the introduction of a common dataset standard, such as the NIfTI standard (https://nifti.nimh.nih.gov/). This was important for the formatting of neuroimaging data. The neuroimaging community gathered together in a consortium to define a standard format for the storage of neuroimaging datasets, the so-called Brain Imaging Data Structure (BIDS) (K. J. Gorgolewski et al., 2016). A common data structure format facilitates the sharing of datasets and makes it possible to create universal neuroimaging toolboxes that work out of the box on any BIDS-conforming dataset. Additionally, through services like OpenNeuro (K. J. Gorgolewski, Esteban, Schaefer, Wandell, & Poldrack, 2017), a free online platform for sharing neuroimaging data, one can test the robustness and flexibility of a new neuroimaging toolbox on hundreds of different datasets.

Software toolboxes like MRIQC (Esteban et al., 2017) and fMRIPrep (Esteban et al., 2019) have shown how fruitful this new neuroimaging ecosystem can be and have highlighted the importance and need for good quality control and high-quality preprocessing workflows with consistent results from diverse datasets. Given the recent developments in the field of data-driven analyses to decode brain states from fMRI time series, there is an increased need for reliable and reproducible statistical analysis of fMRI data, the fundamental input of more advanced machine learning methods, such as Multi-Voxel Pattern Analysis (MVPA) and Convolutional Neuronal Networks (CNNs). Here, we propose an alternative to existing workflows developing to a fully automated pipeline for univariate and multivariate individual and group analyses.

fMRIflows provides flexible temporal and spatial filtering, to account for two recent findings in the data-driven model field. *First*, flexible spatial filtering can become of importance when performing multivariate analysis, as it has been shown that the correct spatial band-pass filtering can improve signal decoding accuracy (Sengupta et al., 2018). *Second*, correct temporal filtering during pre-processing is important and can lead to an improved signal-to-noise ratio (SNR), especially for fMRI datasets with a temporal sampling rate below one second (Viessmann et al., 2018), but only if the filter is applied orthogonally to the other filters during pre-processing to ensure that previously removed noise is not reintroduced into the data (Hallquist et al. 2013; Lindquist et al., 2019). Due to technical improvements in imaging recording through acceleration techniques such as GRAPPA (Griswold et al., 2002) and simultaneous multi-slice/multiband acquisitions (Feinberg et al., 2010; Moeller et al., 2010; Feinberg and Setsompop, 2013), faster sampling rates became possible, to the point that respiratory and cardiac signals can be sufficiently sampled in the BOLD signal. This creates new challenges for the pre-processing of functional images, especially when the external recording of those physiological sources cannot be readily achieved.

fMRIflows presents a consortium of fully automatic neuroimaging pipelines for fMRI analysis, performing standardized pre-processing, as well as 1st- and 2nd-level statistical analyses for univariate and multivariate analysis, with the additional creation of informative quality-control figures. fMRIflows is predicated on the insights and code base of MRIQC (Esteban et al., 2017) and fMRIPrep (Esteban et al., 2019), extending their functionality with regard to the following aspects: (a) flexible temporal and spatial filtering of fMRI data, i.e. low- or band-pass filtering allowing for the exclusion of high-frequency oscillations introduced through physiological noise (Viessmann et al., 2018); (b) accessible and modifiable code base; (c) automatic computation of 1^st^-level contrasts for univariate and multivariate analysis; and (d) automatic computation of 2^nd^-level contrasts for univariate and multivariate analysis.

In this paper, we (1) describe the different pipelines included in fMRIflows and illustrate the different processing steps involved, (2) explain the software structure and setup, and (3) validate fMRIflows’ performance by comparing it to other widely used neuroimaging toolboxes, such as fMRIPrep (Esteban et al., 2019), FSL (Jenkinson et al., 2012) and SPM (Friston et al., 2006).

## 2 Materials and Methods

### 2.1.1 fMRIflows’ processing pipelines

The complete code base of fMRIflows is open access and stored conveniently in six different Jupyter notebooks on https://github.com/miykael/fmriflows. The first notebook does not contain any processing pipeline, but rather serves as a user input document that helps to create JSON files, which will contain the execution-specific parameters for the five processing pipelines contained in fMRIflows: (1) anatomical preprocessing, (2) functional preprocessing, (3) 1^st^-level analysis, (4) 2^nd^-level univariate analysis and (5) 2^nd^-level multivariate analysis. Each of these five pipelines stores its results in a sub-hierarchical folder, specified as an output folder by the user. In the following section, we explain the content of those six Jupyter notebooks.

#### Specification preparation

Each fMRIflows processing pipeline needs specific input parameters to run. Those parameters range from subject ID and number of functional runs per subject to requested voxel resolution after image normalization, etc. Each notebook will read the relevant specification parameters from a predefined JSON file that starts with the prefix “fmriflows_spec”. There is one specification file for the anatomical and functional preprocessing, one for the 1^st^ and 2^nd^ level univariate analysis, and one for the 2^nd^-level multivariate analysis. For an example of these three JSON files, see Supplementary Note 1. The first notebook contained in fMRIflows, called 01_spec_preparation.ipynb, can be used to create those JSON files, based on the provided dataset and some standard default parameters. It does so by using Nibabel v2.3.0 (Brett et al., 2018), PyBIDS v0.8 (Yarkoni et al., 2019) and other standard Python libraries. It is up to the user to change any potential processing parameter should they be different from the used default values.

#### Anatomical preprocessing

The anatomical preprocessing pipeline is contained within the notebook 02_preproc_anat.ipynb and uses the JSON file fmriflows_spec_preproc.json for parameter specification such as voxel resolution. If a specific value is not set, fMRIflows normalizes to an isometric voxel resolution of 1 mm^3^ by default.

However, the user can also choose an anisometric voxel resolution that varies in all three dimensions. Additionally, the user can decide to have a fast or precise normalization. The precise normalization can take up to eight times as long as the fast approach but can provide a more precise alignment. Visual inspection on performed normalization is always desirable since both normalization algorithms may fail in case of a noisy dataset or undetected artifacts. For an example of the JSON file content, see Supplementary Note 1.

The anatomical preprocessing pipeline only depends on the subject-specific T1-weighted (T1w) anatomical images as input files. The individual processing steps are visualized in Figure 1 and consist of: (1) image reorientation, (2) cropping of field of view (FOV), (3) correction of intensity non-uniformity (INU), (4) image segmentation, (5) brain extraction and (6) image normalization. For a more detailed description of the steps involved in this processing pipeline, see Supplementary Note 2.

**Figure 1:**
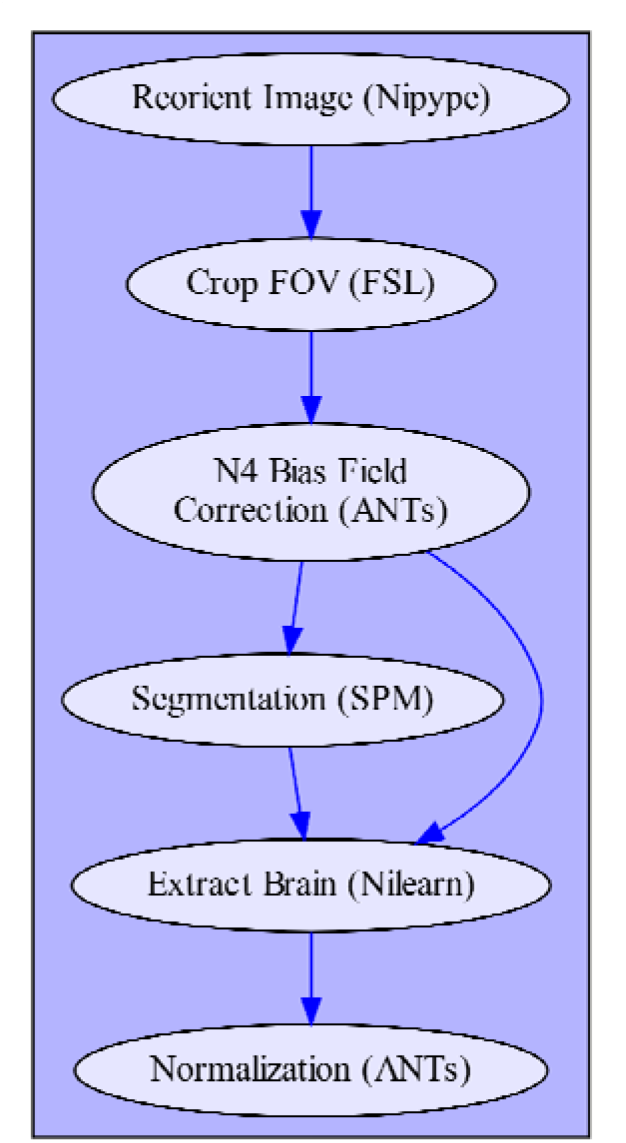
Depiction of fMRIflows’ anatomical preprocessing pipeline. Arrows indicate dependency between the different processing steps and data flow. Name of each node describes functionality, with the corresponding software dependency mentioned in brackets.

#### Functional preprocessing

The functional preprocessing pipeline is contained within the notebook 03_preproc_func.ipynb and uses the JSON file fmriflows_spec_preproc.json for parameter specification. As for specification parameters, users can indicate if slice-time correction should be applied or not, and if so which reference timepoint should be used. The user can also indicate to which isometric or anisometric voxel resolution functional images should be sampled to, and if the sampling is into subject or template space. For the template space, the ICBM 2009c nonlinear asymmetric brain template is used (Fonov et al., 2011). Furthermore, users can specify possible values for low-, high- or band-pass filters in the temporal or spatial domain. Additionally, to investigate nuisance regressors, users can specify the number of CompCor (Behzadi et al., 2007) or independent component analysis (ICA) components they want to extract and which threshold values they want to use to detect outlier volumes. The implications of those parameters will be explained in more detail in the following sections. For an example of the JSON file content, see Supplementary Note 1.

Inputs of the functional preprocessing pipeline depend on the output files from the anatomical preprocessing pipeline, as well as the subject-specific functional images and accompanying descriptive JSON file that contains information about the temporal resolution (TR) and slice order of the functional image recording. This JSON file is part of the BIDS standard and therefore should be available in the BIDS conform dataset. The individual processing steps are schematized in Figure 2 and consist of: (1) image reorientation, (2) non-steady-state detection, (3) creation of functional brain mask, (4) slice time correction, (5) estimation of motion parameters, (6) two-step estimation of coregistration parameters between functional and anatomical image, (7) finalization of motion parameters, (8) single-shot spatial interpolation applying motion correction, coregistration and if specified normalizing images to the template image, (9) construction and application of brain masks, (10) temporal filtering and (11) spatial filtering. It is important to mention that the functional preprocessing is done for each functional run separately to prevent inter-run contaminations. For a more detailed description of the steps involved in this processing pipeline, see Supplementary Note 3.

**Figure 2:**
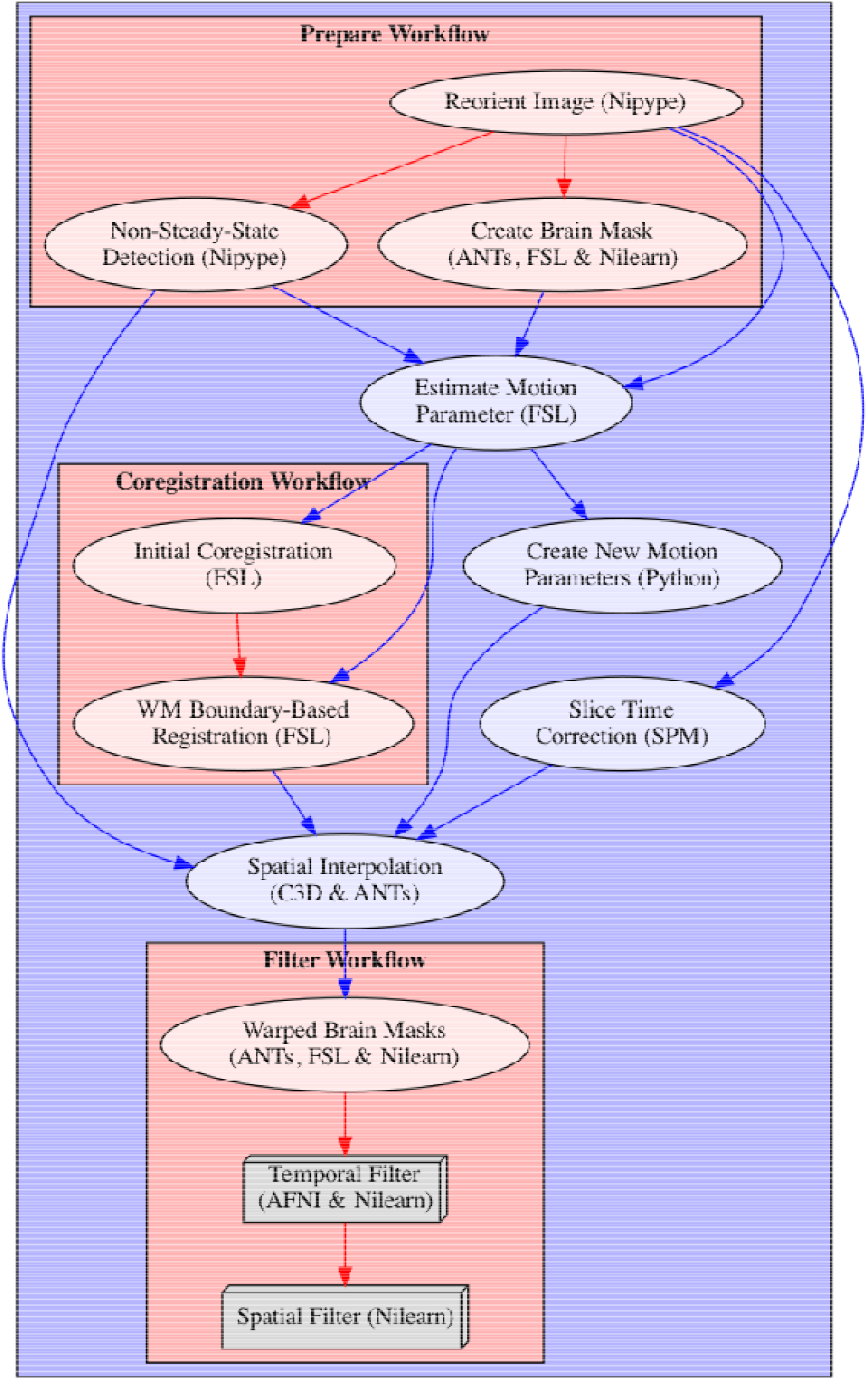
Depiction of fMRIflows’ functional preprocessing pipeline. Arrows indicate dependency between the different processing steps and data flow. The name of each node describes the functionality, with the corresponding software dependency mentioned in brackets. Steps that can be grouped into specific sections are contained within a red box to facilitate understanding of the pipeline. The color of the arrows indicates if the connection stays within a section (red) or not (blue). Nodes depicted as gray boxes indicate that they can be run multiple times with iterating input values, i.e. performing a spatial smoothing with an FWHM value of 4 and 8mm.

#### 1st-level analysis

The first-level analysis pipeline is contained within the notebook 04_analysis_1st-level.ipynb and uses the JSON file fmriflows_spec_analysis.json for parameter specification. As for specification parameters, users can indicate which nuisance regressors to include in the GLM, if outliers should be considered, and if the data is already in template space or if this normalization should be done after the estimation of the contrasts. Users can also specify other GLM model parameters, such as the high-pass filter value and the type of basis function that should be used to model the hemodynamic response function (HRF). Additionally, the users will also specify a list of contrasts they want to be estimated, or if they want to create specific contrasts for each stimulus column in the design matrix, and/or for each session separately, which then later might also be used for multivariate analysis. For an example of the JSON file content, see Supplementary Note 1.

The 1st-level analysis pipeline depends on a number of outputs from the previous anatomical and functional preprocessing pipelines, i.e. the TSV (tab separated value) file containing motion parameters and confound regressors, a text file indicating the number of non-steady-state volumes removed from the functional image, and a text file containing a list of indexes identifying outlier volumes. Additionally, the 1st-level analysis pipeline also requires BIDS conform events files containing information on the applied experimental design, including types of conditions and their respective onsets and durations. The individual processing steps included in the 1st-level analysis consist of: (1) collecting preprocessed files and model relevant information, (2) model specification and estimation, (3) univariate contrast estimation, (4) optional preparation for multivariate analysis, (5) optional spatial normalization of contrasts (Figure 3). All of the relevant steps, that is model creation, estimation and contrast computation are performed with SPM version 12. For a more detailed description of the steps involved in this processing pipeline, see Supplementary Note 4.

**Figure 3:**
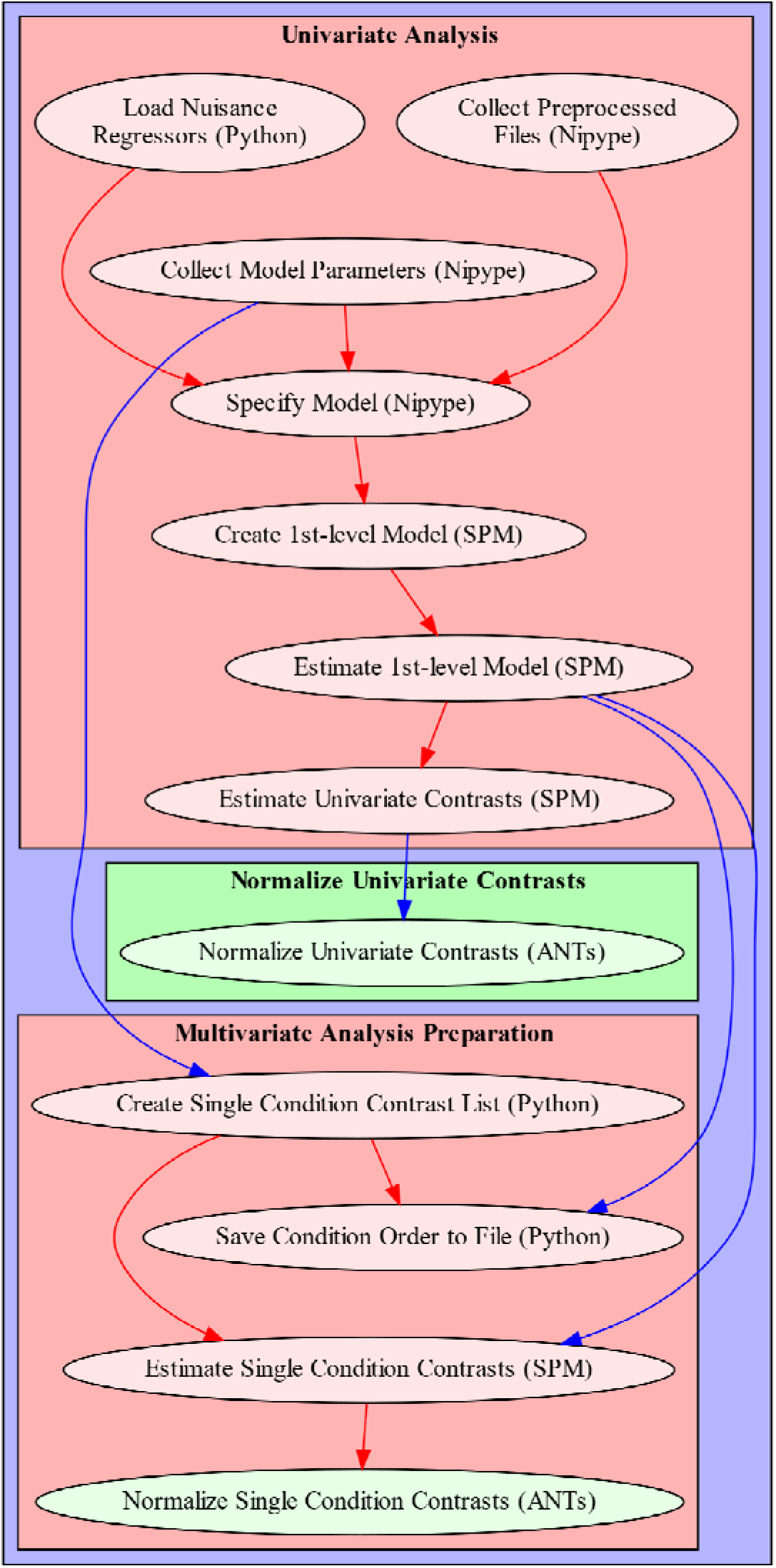
Depiction of fMRIflows’ 1st-level analysis pipeline. Arrows indicate dependency between the different processing steps and data flow. The name of each node describes the functionality, with the corresponding software dependency mentioned in brackets. Sections that can be grouped into specific sections are contained within a red box to facilitate understanding of the pipeline. The color of the arrows indicates if the connection stays within a section (red) or not (blue). Nodes depicted in green are optional and can be run if spatial normalization was not yet performed during preprocessing.

#### 2nd-level univariate analysis

The second-level univariate analysis pipeline is contained within the notebook 05_analysis_2nd-level.ipynb and uses the JSON file fmriflows_spec_analysis.json for parameter specification. Users can specify the probability value used as a cutoff for the threshold of the GM probability tissue map in template space that is later used during the model estimation. Additionally, users can specify voxel- and cluster-threshold topological thresholding of the statistical contrast, as well as relevant AtlasReader (Notter et al., 2019) parameters for the creation of the output tables and figures.

The 2nd-level univariate analysis pipeline depends only on the estimated contrasts from the 1st-level univariate analysis. No further contrast specification is required as fMRIflows currently only implements a simple one-sample T-test. The individual processing steps included in the 2nd-level univariate analysis consist of: (1) gathering of the 1st-level contrasts, (2) creation and estimation of the 2nd-level model, (3) estimation of contrast estimation, (4) topological thresholding of contrasts, (5) results creation with AtlasReader. As for the 1st-level analysis, all of the relevant model creation, estimation and contrast computations are performed with SPM12. For a more detailed description of the steps involved in this processing pipeline, see Supplementary Note 5.

#### 2nd-level multivariate analysis

The second-level multivariate analysis pipeline is contained within the notebook 06_analysis_multivariate.ipynb and uses the JSON file fmriflows_spec_multivariate.json for parameter specification. Users can define a list of classifiers to use for the multivariate analysis, the sphere radius and the step size of the searchlight approach. To perform a 2nd-level analysis of searchlight results users can decide between a classical GLM approach testing against chance level and a more recommended permutation-based method as described in Stelzer, Chen, & Turner (2013) with the option of determining the number of permutations. Additionally, users can specify voxel- and cluster-threshold topological thresholding of the statistical contrast, as well as relevant AtlasReader parameters for the creation of the output tables and figures.

The 2nd-level multivariate analysis pipeline depends on the estimated contrasts from the 1st-level multivariate analysis, the associated CSV file containing a list of the corresponding contrast labels and a list of binary classification identifiers. In contrast to the other notebooks, this notebook uses Python 2.7 to accommodate the requirements of PyMVPA v2.6.5 (Hanke et al., 2009). The individual processing steps included in the 2nd-level multivariate analysis consist of: (1) data preparation for the analysis with PyMVPA, (2) searchlight classification, (3) computation of group analysis using a T-test, (4) computation of group analysis according to Stelzer et al. (2013), and (5) results creation with AtlasReader. For a more detailed description of the steps involved in this processing pipeline, see Supplementary Note 6.

### 2.1.2 Infrastructure and access to fMRIflows

The source code of fMRIflows is available at GitHub (https://github.com/miykael/fmriflows) and is licensed under the BSD 3-Clause “New” or “Revised” License. The code is written in Python v3.7.2 (https://www.python.org), stored in Jupyter Notebooks v4.4.0 (Kluyver et al., 2016) and distributed via Docker v18.09.2 (https://docker.com) containers that are publicly available via Docker Hub (https://hub.docker.com). The usage of Docker allows the user to run fMRIflows on any major operating system, with the following command:

docker run -it -p 9999:8888 -v /home/user/ds001:/data miykael/fmriflows

The first flag -it indicates that the docker container should be run in interactive mode, while the second flag -p 9999:8888 defines the port (here 9999) that we want to use to access the Jupyter Notebooks via the web-browser. The third flag, -v /home/user/ds001:/data tells fMRIflows the location of the BIDS-confirm dataset that should be mounted in the docker container, here located at /home/user/ds001. Once the docker container is launched, the interactive Jupyter Notebooks can be accessed through the web browser. Further detailed documentation can be found in the github repository.

fMRIflows uses many different software packages for the individual processing steps. Specifically, this entails the following: Nipype v1.1.9 (Gorgolewski et al., 2011), FSL v5.0.9 (Smith et al., 2004), ANTs v2.2.0 (Avants et al., 2011), SPM12 v7219 (Penny et al., 2011), AFNI v18.0.5 (Cox and Hyde, 1997), Nilearn v0.5 (Abraham et al., 2014), Nibabel v2.3.0 (Brett et al., 2018), PyMVPA v2.6.5 (Hanke et al., 2009), Convert3D v1.1 (https://sourceforge.net/p/c3d), AtlasReader v0.1 (Notter et al., 2019) and PyBIDS v0.8 (Yarkoni et al., 2019). In addition to some standard Python libraries, fMRIflows also uses Numpy (Oliphant, 2007), Scipy (Jones et al., 2001), Matplotlib (Hunter, 2007), Pandas (McKinney and others, 2010) and Seaborn (http://seaborn.pydata.org).

With every new pull request pushed to the GitHub repository of fMRIflows, a test instance on CircleCI (https://circleci.com) is deployed to test the complete code base for execution errors. This framework allows the continuous integration of new code to fMRIflows, and guarantees the general functionality of the software package. Outputs are not controlled for their correctness.

### 2.1.3 Validation of fMRIflows

fMRIflows was validated in two phases. In *Phase 1*, we validated the proficiency of the toolbox by applying it on different kinds of fMRI datasets conforming to the BIDS standard (K. J. Gorgolewski et al., 2016) available via OpenNeuro.org (Gorgolewski et al., 2017). Insights during this phase allowed us to improve the code base and make fMRIflows robust to a diverse set of datasets. In *Phase 2*, we compared the performance of the toolbox to similar neuroimaging preprocessing pipelines such as fMRIPrep, FSL, and SPM. To better understand where fMRIflows overlaps or diverges from comparable processing pipelines, we investigated the preprocessing, subject-level and group-level outcomes for all four toolboxes, run on three different datasets.

#### Phase 1: Proficiency validation

To investigate the capabilities and flaws of the initial implementation of the toolbox, fMRIflows was run on different datasets, either available publicly via OpenNeuro.org or available privately to the authors. Such an approach allowed the exploration of datasets with different temporal and spatial resolutions, SNRs, FOVs, numbers of slices, scanner characteristics, and other sequence parameters, such as acceleration factors and flip angles.

#### Phase 2: Performance validation

To validate the performance of fMRIflows, we used three different task-based fMRI datasets and compared its preprocessing to the three neuroimaging processing pipelines/software packages fMRIPrep, FSL and SPM. The comparison was done on preprocessing, subject-level and group-level outputs. Because of differences in how FSL and SPM perform subject- and group-level analyses and due to the lack of such routines in fMRIPrep, all subject- and group-level analyses for the performance validation were performed using identical Nistats routines.

The three datasets (see Table 1) were all acquired on scanners with a magnetic field strength of 3 Tesla and differ in many sequence parameters, most notably in the temporal resolution with which they were recorded. This is especially important as we aim to highlight that the right handling of temporal filtering is crucial for datasets with a temporal resolution below 1000ms.

**Table 1:**
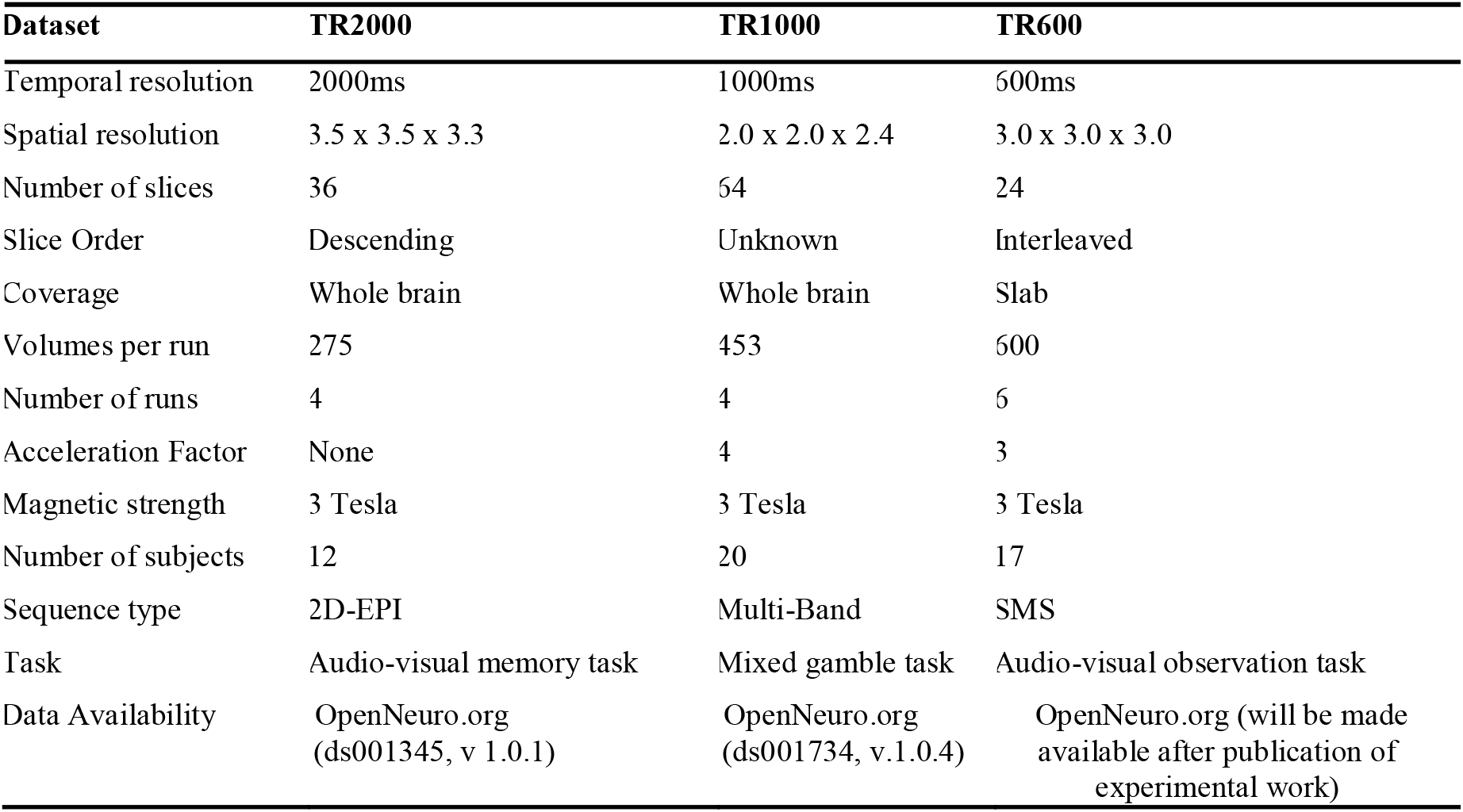
Overview of the datasets used to validate fMRIflows.

**Dataset TR2000** has a comparably low temporal sampling and spatial resolution. It serves as a standard dataset, recorded with a standard EPI scan sequence. The dataset and paradigm are described in more detail in Notter et al. (under review). In short, participants performed a continuous recognition task and indicated for each image whether it is old or new. When the image was presented for the first time (new) it was either presented with no sound (unisensory visual context) or together with a sound (multisensory context).

**Dataset TR1000** has a rather high temporal sampling and spatial resolution and serves as an advanced dataset, recorded with a scan sequence using a multiband acceleration technique. The dataset and paradigm are described in more detail in Botvinik-Nezer et al. (2019). In short, participants performed a mixed gambling task in which they were asked to either accept or reject a possible monetary gain or loss.

**Dataset TR600** has a very high temporal sampling with a moderate spatial resolution and serves as an extreme dataset, recorded with scan sequences using a simultaneous multi-slice (SMS) acceleration technique (Feinberg et al., 2010). This dataset was collected for another project. In short, participants were shown auditory, visual or audiovisual stimuli containing either an animal (as an image or sound), pure noise or both together. Participants performed a discrimination task in which they had to indicate if they perceived a stimulus with an animal in it or not, independent of the stimuli modality. The stimuli were either presented in a unisensory or multisensory context.

All participants of the Datasets TR2000 and TR600 have been included in the performance validation, while only the first 20 out of the 120 total participants of the Dataset TR1000 were used in order to reduce computation time and make this dataset comparable to the other two. Datasets TR2000 and TR1000 are already publicly available through the OpenNeuro platform. Dataset TR600 is in preparation to be published on OpenNeuro as well. Until then, this dataset is available upon request to the corresponding author.

The **pre-processing pipelines** with fMRIflows, fMRIPrep, FSL and SPM were based on the default parameters and only differed in the following points from their standard implementations: (1) Functional images were resampled to an isometric voxel resolution according to the dominant resolution dimension within a dataset; (2) Spatial smoothing of the functional images is applied after preprocessing of the images, using a Nilearn routine and a smoothing kernel with a Full Width at Half Maximum (FWHM) of 6mm, in order to keep the approaches comparable, as spatial smoothing is not included in the fMRIPrep workflow; (3) Anatomical images in the FSL pipeline were first cropped to a standard FOV, followed by brain extraction using FSL’s BET before FSL’s FEAT was launched; (4) In the case of FSL, the normalization from structural to standard space was done using a non-linear warping approach with 12 degrees of freedom and a spline interpolation model; (5) In the case of SPM, the template brain for the normalization was its standard tissue probability brain TPM, while for fMRIflows, fMRIPrep and FSL, the ICBM 2009c nonlinear asymmetric brain template was used.

The statistical inference was not performed on any of the investigated toolboxes to prevent the introduction of a software-specific bias. The 1^st^- and 2^nd^-level analysis was performed using Nistats, Nilearn and other Python toolboxes and only differed between the toolboxes in the following ways: (1) the estimated motion parameters added to the design matrix during the 1^st^-level analysis differed for each toolbox as they were based on the software-specific preprocessing routine; (2) the number of volumes per run used during the 1^st^-level analysis of fMRIflows might differ slightly from the other approaches, as the fMRIflows routine removes non-steady state volumes during the preprocessing; (3) SPM used its own tissue probability map to create a binary mask restricted to gray matter voxels during the group analysis, while the other three toolboxes used the ICBM 2009c gray matter probability map instead.

To compare the unthresholded group statistic maps between the toolboxes, we created for each pairwise combination of preprocessing approach a Bland-Altman 2D histogram plot, as described by (Bowring, Maumet, & Nichols, 2018). These plots show the difference between the statistic value (y-axis), against the mean statistic value (x-axis) for all voxels within the intersection of the respective brain mask. In other words, it summarized in a 2D histogram plot, for each voxel how much higher the statistical value in toolbox B is (y-axis), in comparison to toolbox A’s statistical value (x-axis).

The complete lists of parameters, the scripts to perform preprocessing, 1^st^- and 2^nd^-level analysis and the scripts to create individual figures can be found on the fMRIflows GitHub page (https://github.com/miykael/fmriflows/tree/master/paper). Derivatives generated for the validation in phase 2 can be inspected and downloaded on NeuroVault (Gorgolewski et al., 2015) under the following links: (1) Standard deviation maps of temporal averages after preprocessing (https://identifiers.org/neurovault.collection:5645), (2) temporal SNR maps after preprocessing (https://identifiers.org/neurovault.collection:5713), (3) binarized 1st-level activation count maps (https://identifiers.org/neurovault.collection:5647), (4) 2nd-level activation maps (https://identifiers.org/neurovault.collection:5646).

## 2.2 Results

### 2.2.1 Summary of outputs obtained by fMRIflows’ processing pipelines

#### Output generated after executing the anatomical preprocessing pipeline

After the execution of the anatomical preprocessing pipeline, the following files are generated for each subject: (1) image of the inhomogeneity-corrected full head image, (2) image of the extracted brain, (3) binary mask used for the brain extraction, (4) individual tissue probability maps for gray matter (GM), white matter (WM), cerebrospinal fluid (CSF), skull and head, (5) normalized anatomical image in template space (6) reverse-normalized template image in subject space, (7) plus the corresponding transformation matrices used for output 5 and 6. Each anatomical preprocessing output folder also contains (8) the ICBM 2009c brain template used for the normalization, sampled to the requested voxel resolution.

In addition to these files, the following three informative figures are generated: (1) tissue segmentation, (2) brain extraction and (3) spatial normalization of the anatomical image. A shortened version of those three figures, as well as their explanation, are shown in Figure 4.

**Figure 4:**
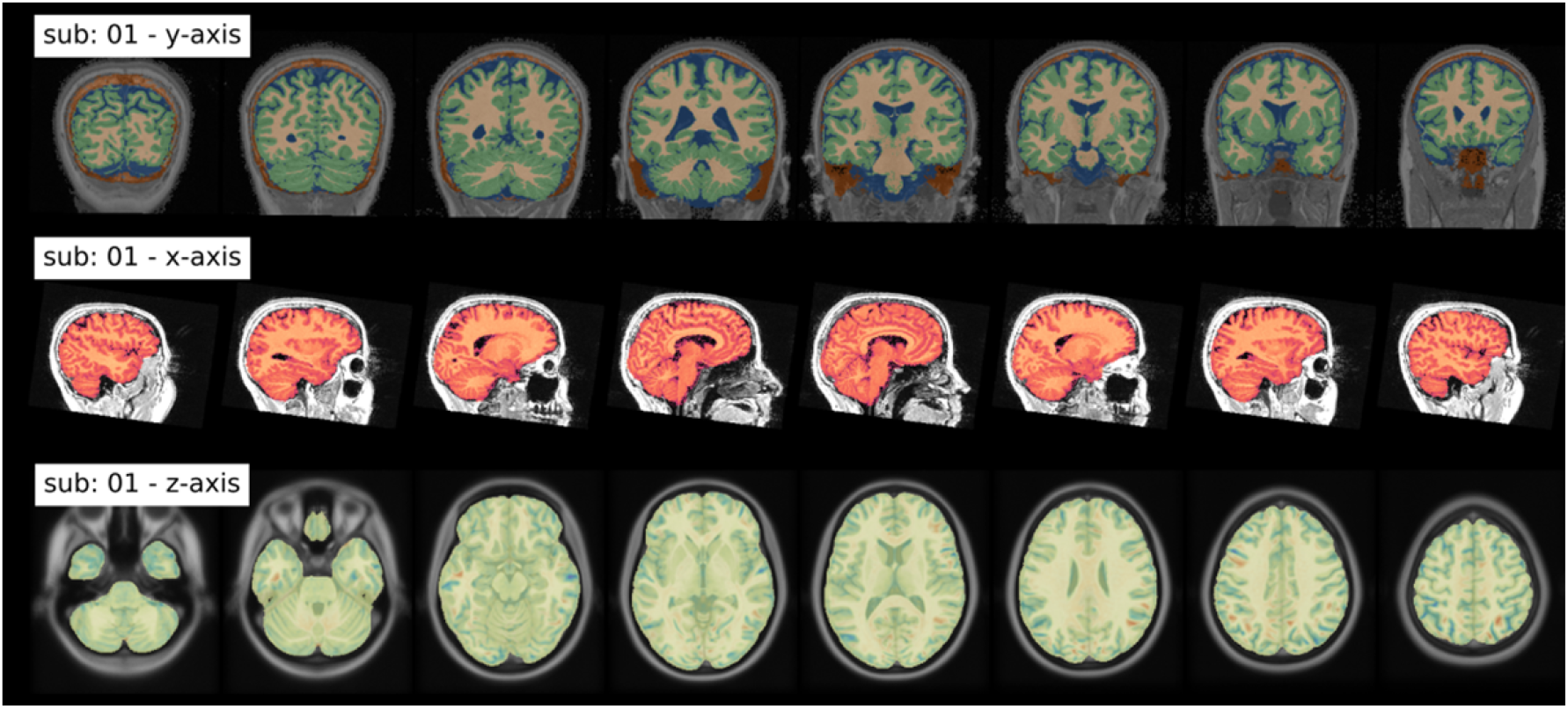
Summary of output figures generated by fMRIflows after executing the anatomical preprocessing pipeline. (***Top***) Coronal view of the image segmentation output, showing gray matter tissue in green, white matter tissue in beige, and cerebrospinal fluid in blue. (***Middle***) Sagittal view of the brain extraction output, showing the extracted brain image in red, and the original anatomical image in gray. (***Bottom***) Axial view of the spatial normalization output, showing the normalized brain image highlighted in yellow, overlaid over the ICBM 2009c brain template in gray. Regions in red and blue show negative and positive deformation discrepancies between the normalized subject image and the template.

#### Output generated after executing the functional preprocessing pipeline

After the execution of the functional preprocessing pipeline, the following files are generated separately for each subject, each functional run and each temporal filtering: (1) text file indicating which volumes were detected as outliers, (2) tabular separated (TSV) file containing all extracted confound regressors, (3) text file containing the six motion parameter regressors according to FSL’s output scheme, (4) binary masks for the brain, (5) masks for anatomical and functional component based noise correction, (6) functional mean image, and (7) completely preprocessed functional images, separated by spatial smoothing approaches. Each subject folder also contains (8) one text file per functional run indicating the number of non-steady-state volumes at the beginning of the run.

The following is a more detailed description of the multiple confounds fMRIflows estimates during functional preprocessing:

##### Confounds based on motion parameters

In addition to the head motion parameters created during preprocessing, fMRIflows also computes (1) 24-parameter Volterra expansion of the motion parameters (Friston et a., 1996) using custom scripts and (2) Framewise Displacement (FD) component (Power et al., 2012) using Nipype.

##### Confounds based on global signal

Functional images before spatial smoothing were used to compute confound regressors, such as (1) DVARS, which represents the spatial standard deviation of the signal after temporal differencing, to identify motion-affected frames (Power et al., 2012), using Nipype and (2) four global signal curves representing the average signal in the total brain volume (TV), GM, WM and CSF, using Nilearn.

##### Detection of outlier volumes

The user can specify which of the six signal curves for FD, DVARS and average signal in TV, GM, WM and CSF to use to identify outlier volumes (see Figure 5A). Those are volumes that have larger fluctuations in the signal values in a given volume, compared to the z-scored standard deviation throughout the time course. The exact threshold for each curve can be adapted by the user, but its default value is set to a z-value of 3.27, representing 99%, for the FD, DVARS and TV signal. The identification number of each outlier volume is stored in a text file that might be used in the 1^st^-level pipeline during the GLM model estimation to remove the effect of those volumes from the overall analysis, also known as censoring (Caballero-Gaudes and Reynolds, 2016).

**Figure 5:**
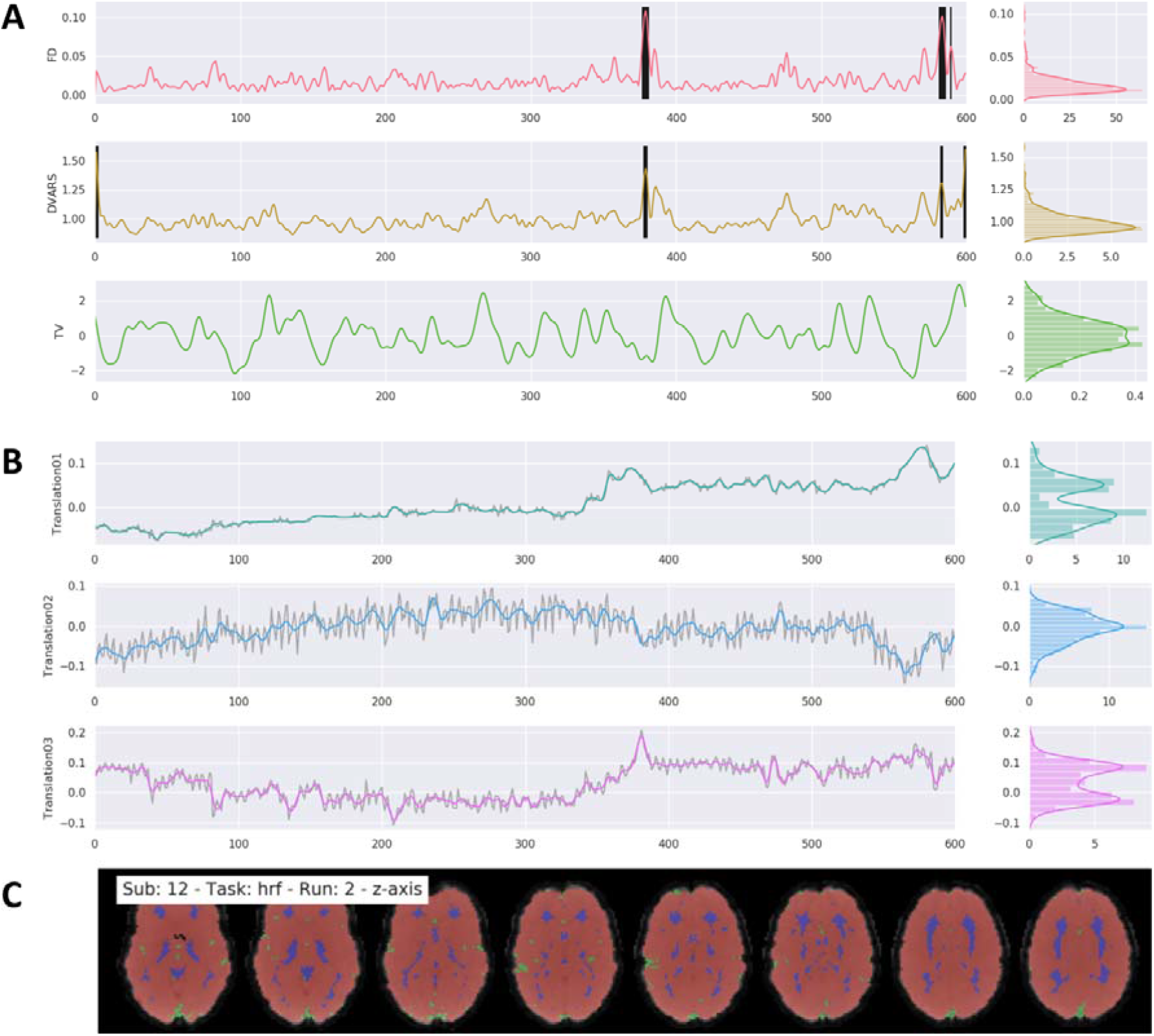
Example of general output figures generated by fMRIflows after executing the functional preprocessing pipeline. The dataset used to generate these figures was recorded with a TR of 600ms and had a total of 600 volumes per run. Preprocessing included a low-pass filter at 0.2 Hz. Distribution plots on the right side of the figures in parts A and B represent value frequency in y-direction. **A)** Depiction of the nuisance confounds FD, DVARS and TV. Detected outlier volumes are highlighted with vertical black bars. **B)** Estimation of translation head motion after application of low-pass filtering at 0.2 Hz in color, and before temporal filtering in light gray. **C)** Depiction of brain masks used to compute DVARS (red), temporal (green) and anatomical (blue) CompCor confounds, overlaid on the mean functional image (grey).

##### Confounds based on signal components

Using the temporal filtered functional images, two different kinds of approaches are performed to extract components that could be used for denoising or dimensionality reduction of the data. The first approach is called CompCor (Behzadi et al., 2007) and uses principal component analysis (PCA) to estimate the main sources of noise within specific confound regions. Regions are either defined by their temporal or anatomical characteristics. The temporal CompCor approach (tCompCor) considers the 2% most variable voxels within the confound brain mask as sources of confounds. The anatomical CompCor approach (aCompCor), considers voxels within twice eroded WM and CSF brain masks as sources of confounds. The user can specify how many aCompCor and tCompCor components should be computed, but the default value is set to five each. The second approach uses independent component analysis (ICA) to perform source separation in the signal (Figure 6). Using Nilearn’s CanICA routine, fMRIflows computes by default the top ten independent components throughout the confound masks. The number of confounds to extract can be adjusted by the user. It is the user’s responsibility to evaluate appropriately whether residual artifacts are present and need to be removed.

**Figure 6:**
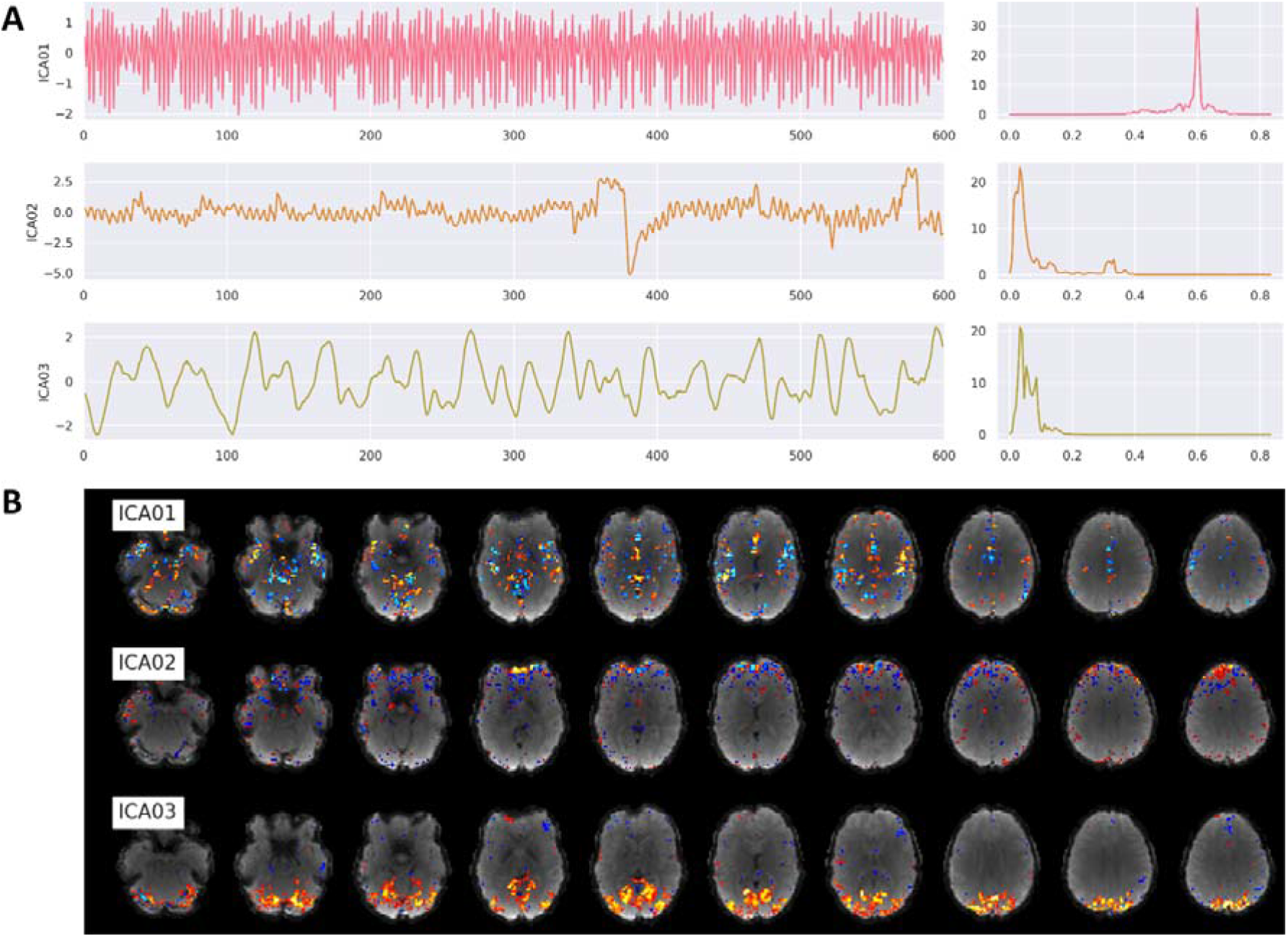
Example of ICA output figures generated by fMRIflows after executing the functional preprocessing pipeline. The dataset used to generate these figures was recorded with a TR of 600ms and had a total of 600 volumes per run. **A)** Correlation between the first three ICA components and the functional image over time (left) and the corresponding power density spectrum with frequency on the x-axis (right). The first component most likely depicts respiration at 0.6 Hz, while the third component is most likely visual activation induced by the visual stimulation task during data acquisition. B) Correlation strength between a given ICA component and the location in the brain volume for the first three ICA components.

##### Storage of confound information

All of the confound curves computed after functional preprocessing are stored in a TSV file to allow for easy access.

##### Diverse set of overview figures

To allow for visual inspection of the numerous outputs generated after the execution of the functional preprocessing pipeline, fMRIflows creates many informative overview figures. These overviews cover the motion parameters used for head motion correction, the anatomical and temporal CompCor components, FD, DVARS, the average signal in TV, GM, WM and CSF, and the ICA components. fMRIflows also creates a brain overview figure showing the extent of the different masks applied during functional preprocessing, a spatial correlation map between the ICA components and the individual voxel signal, and a carpet plot according to (Power, 2017) and (Esteban et al., 2019). To better visualize underlying structures in the carpet plot the time series traces are sorted by their correlation coefficients to the average signal within a given region, allowing for a positive or negative time lag of 2 volumes. A shortened version of all these figures, as well as their explanations, are shown in Figures 5-7.

**Figure 7:**
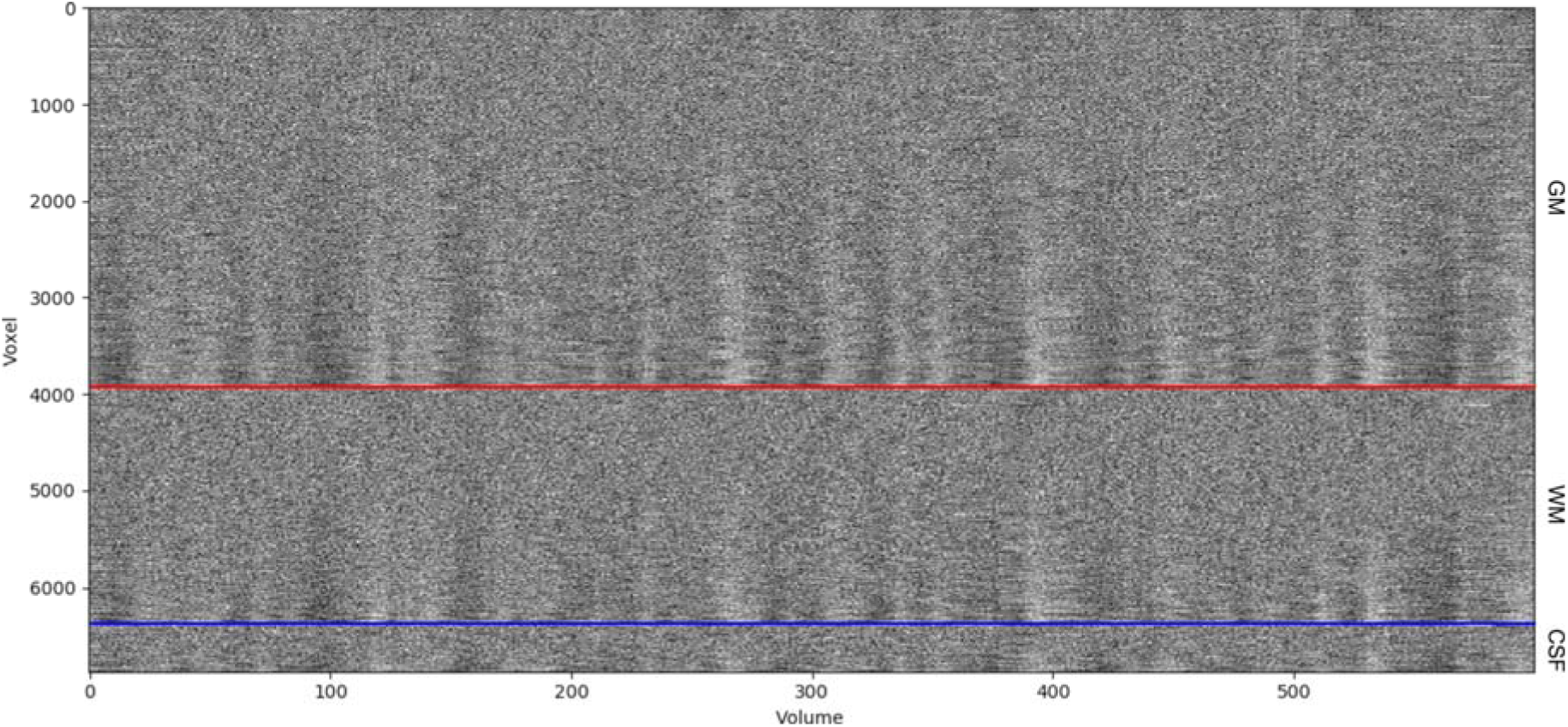
Example of a carpet plot figure generated by fMRIflows after executing the functional preprocessing pipeline. The dataset used to generate this figure was recorded with a TR of 600ms and had a total of 600 volumes per run. This panel shows the signal after preprocessing for every other voxel (y-axis), over time in volumes (x-axis). The panel shows voxels in the gray matter (top part), white matter (between blue and red line) and CSF (bottom section). The data was standardized to the average signal and ordered within a given region according to the correlation coefficient between a voxel and the average signal of this region.

#### Output generated after executing the 1st-level analysis pipeline

After the execution of the 1st-level analysis, the following files are generated for the univariate analysis: (1) contrasts and statistical map of the specified contrasts, (2) SPM.mat file containing the information relevant for the model, (3) visualization of the design matrix used in the 1st-level model depicting the regressor for the stimuli, motion and confounds, and (4) glass brain plot for each estimated contrast thresholded at the top 2% of positive and negative values created with AtlasReader to provide a general overview of the quality of contrasts. The multivariate analysis part of this notebook creates: (1) one contrast image per condition and session which later can be used as samples for the multivariate analysis, and (2) a label file identifying the condition of each contrast.

#### Output generated after executing the 2nd-level analysis pipeline

After the execution of the 2nd-level univariate analysis, the following files are generated, individually for each contrast and spatial and temporal filter that was applied: (1) contrasts and statistical map of one-sample *t*-test contrast, (2) SPM.mat file containing the information relevant for the model, (3) thresholded statistical maps with corresponding AtlasReader outputs (i.e. glass brain plot to provide a result overview, cross section plot showing each significant cluster individually, informative tables concerning the peak and cluster extent of each cluster).

After the execution of the 2nd-level multivariate analysis, the following files are generated, for each specified comparison individually: (1) subject-specific permutation files needed for correction according to (Stelzer et al., 2013), (2) group-average prediction accuracy maps as well as corresponding feature-wise maps representing chance level acquired via a bootstrapping approach (Stelzer et al., 2013), (3) group-average prediction accuracy maps after correction for multiple comparisons and (4) thresholded statistical result maps with corresponding AtlasReader outputs (i.e. glass brain plot to provide a result overview, cross section plot showing each significant cluster individually, informative tables concerning the peak and cluster extent of each cluster).

### 2.2.2 Results of phase 1: Proficiency validation

Due to differences in scanner hardware, scan protocols, research requirements and expertise of the person who records the images, fMRI datasets can come in many different shapes and forms. We ran fMRIflows on several datasets to make sure that it is capable of dealing with differences inherent to each of them. In this section, we summarize the main issues we encountered during this process and describe how we tackled each of them.

#### Image orientation

fMRIflows reorients all anatomical and functional images at the beginning of the preprocessing pipeline to the neurological convention RAS (right, anterior, superior) to prevent failures of coregistration between anatomical and functional images due to orientation mismatches within subjects.

#### Image extent

Some datasets have unusually large image coverage along the inferior-superior axis, which means that their anatomical images also often contain part of the participant’s neck. This can lead to unwanted outcomes in certain neuroimaging routines, as they were not tested for such additional tissue coverage. This is most pronounced in the case of FSL’s BET routine, which has difficulty finding the center and extent of the brain, or SPM’s segmentation routine which depends on the distribution of the voxel intensities within the whole volume. To prevent these and other unforeseen behaviors, fMRIflows uses FSL’s robustfov routine to restrict all anatomical images to the same spatial extent.

#### Image inhomogeneity

Depending on the scan sequence protocol or the scanner hardware itself, some datasets can contain strong image intensity inhomogeneities, caused by an inhomogeneous bias field during data acquisition. This can have a negative effect on many different neuroimaging routines, most pronounced in brain extraction and image segmentation. To tackle this issue, fMRIflows uses ANTs’ N4BiasFieldCorrection routine, which allows the analysis of datasets with even low image quality and strong image inhomogeneity. In the anatomical preprocessing pipeline, inhomogeneity correction is applied to improve the final output image. In the functional preprocessing pipeline, inhomogeneity correction is only applied to improve the estimation and extraction of different tissue types but does not directly change the values in the final output image.

#### Brain extraction

Different brain extraction routines were explored to ensure: 1) that the extraction is sufficiently robust to handle different kinds of datasets, 2) that it is neither too conservative nor liberal with the removal of non-brain tissues, and 3) that it has an overall reasonably fast computation time. The best and most consistent results were achieved using SPM’s image segmentation routine, followed by specific thresholding and merging of the GM, WM and CSF probability maps. FSL’s BET routine was not robust enough to lead to stable results on all tested datasets. While ANTs’ Atropos routine led to comparably good results, we went with SPM because of the much faster computation time.

#### Image interpolation

For the single-shot spatial interpolation during normalization, we used ANTs and explored NearestNeighbor, BSpline and LanczosWindowedSinc (Lanczos, 1964) interpolation. NearestNeighbor interpolation led to unnatural-looking voxel-to-voxel value transitions. BSpline led in general to good results, but had issues, especially with datasets that did not have full brain coverage and introduced some rippling low-value fluctuations at the borders of non-zero voxels. LanczosWindowedSinc interpolation led to the best outcome by minimizing the smoothing effects and preventing the introduction of additional confounds reaffirming the observations from fMRIPrep (Esteban et al., 2019).

### 2.2.3 Results of phase 2: Performance validation

The performance validation of fMRIflows was conducted on three different task-based fMRI datasets, as described in Table 1. The preprocessing of fMRIflows was compared to other neuroimaging processing pipelines such as fMRIPrep, FSL and SPM. We tested fMRIflows’ preprocessing pipeline with and without a temporal low-pass filter of 0.2 Hz to better understand performance differences between toolboxes and to stress the importance of adequate temporal filtering when processing fMRI datasets with high temporal resolution.

#### Estimated spatial smoothness after functional preprocessing

Each preprocessing step that resamples a functional image, such as slice time correction, motion correction, and spatial or temporal interpolation has the potential to increase the spatial smoothness in the data. The less smoothness is introduced during preprocessing, the closer the data are to their initial version. We used AFNI’s 3dFWHMx to estimate the average spatial smoothness (FWHM) of each functional image after preprocessing to compare the amount of data manipulation that was applied to the raw data (see Figure 8). As this FWHM value depends on the voxel resolution of a given dataset, we normalized it by the volume of the voxel to achieve a common FWHM value per 1mm^3^.

**Figure 8:**
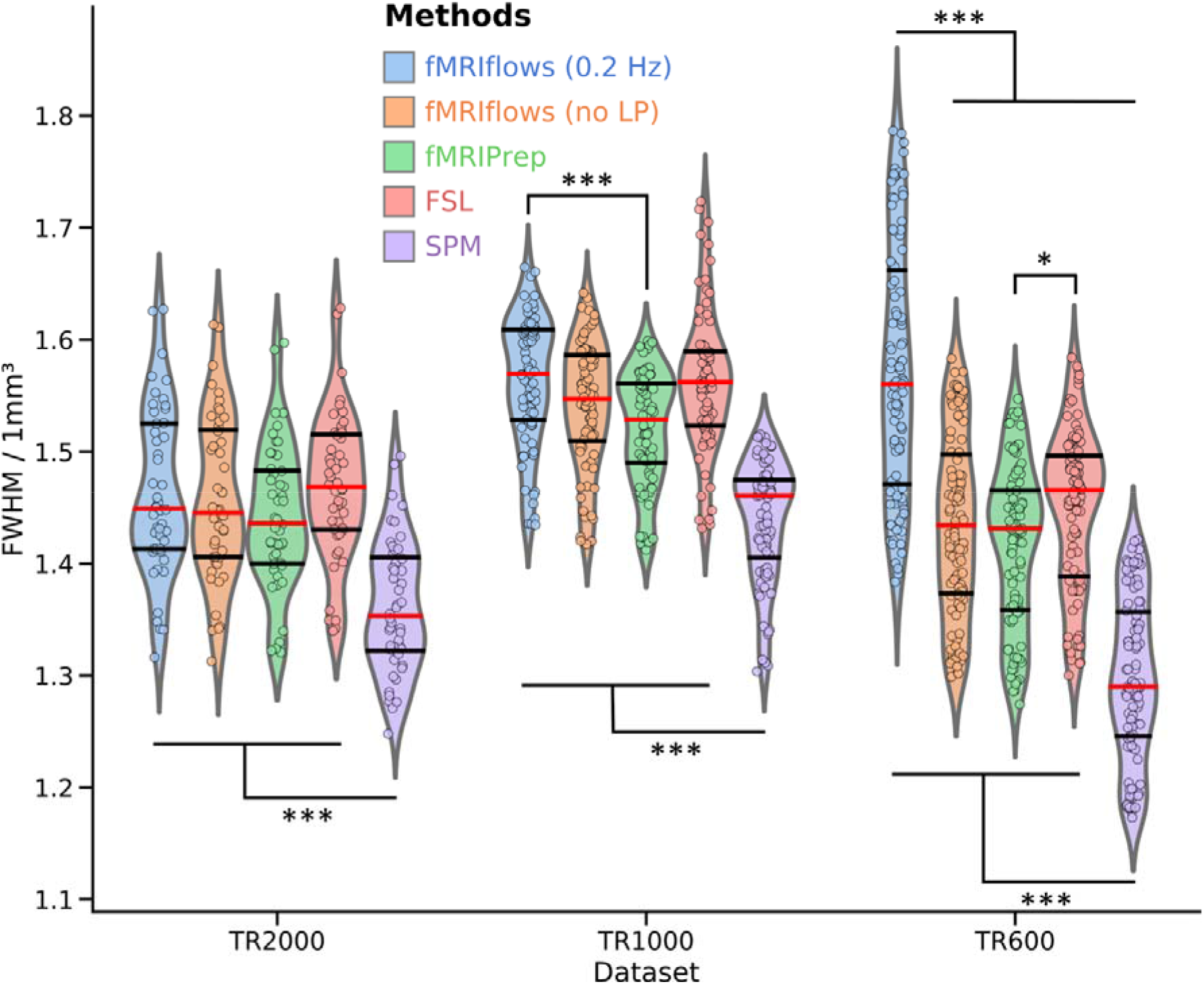
Investigation of estimated spatial smoothness after functional preprocessing of three different datasets, processed with varying approaches. The five different preprocessing approaches fMRIflows with (blue) and without (orange) a low-pass filter at 0.2 Hz, fMRIPrep (green), FSL (red) and SPM (violet) are plotted separately for the dataset TR2000 (left), TR1000 (middle) and TR600 (right). The violin plots indicate the overall distribution of the normalized smoothness estimates of each functional image (depicted in individual dots: TR2000=48 dots, TR1000=80 dots, TR600=102 dots). The red horizontal line represents the median value, while the horizontal black lines indicate the 25 and 75 percentiles of the value distribution respectively. Two-sided t-tests were computed for each pair of approaches used and each dataset. Significant differences between groups are indicated with *: *p*<0.05 and ***: *p*<0.001.

Overall, the estimated spatial smoothness after preprocessing with fMRIflows (without a low-pass filter) is comparable to the one with fMRIPrep, while SPM’s is in general significantly lower and FSL’s is slightly higher. The differences with respect to SPM are probably due to the fewer numbers of resampling steps involved in SPM’s preprocessing pipeline. The differences with respect to FSL are probably due to the interpolation method used during image resampling. While the FSL preprocessing pipeline uses the spline interpolation, fMRIflows and fMRIPrep use the LanczosWindowedSinc interpolation, which is known to minimize the smoothing during interpolation. The application of a temporal low-pass filter at 0.2 Hz during fMRIflows’ preprocessing leads to a significantly higher spatial smoothness for the TR600 dataset when compared with the other approaches. This effect might also be present in the TR1000 dataset. However, there the difference between the fMRIflows preprocessing with and without low-pass filtering is not significant. This increased spatial smoothness for the approach that uses a low-pass filter makes sense, as the goal of the temporal low-pass filter itself is to smooth the time series values. This temporal smoothing forcibly also increases the spatial smoothness at each individual time point.

#### Performance check of spatial normalization

We computed the standard deviation map for each population, based on the temporal average map of each preprocessed functional image, to compare the performance of spatial normalization of the different preprocessing methods on the three different datasets (see Figure 9).

**Figure 9:**
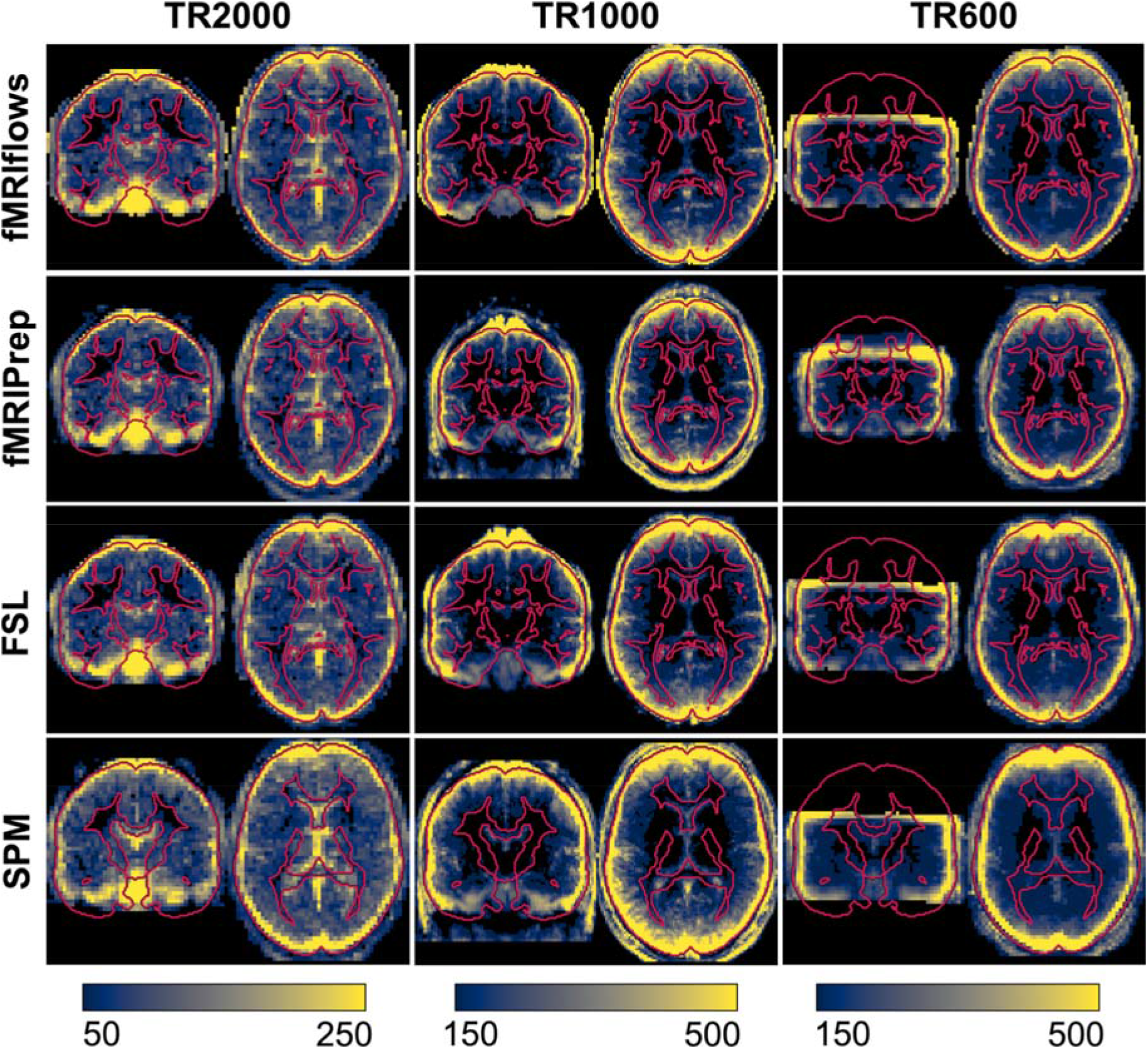
Depiction of standard deviation maps of the temporal averages of three different datasets, after multiple functional preprocessing approaches. Preprocessing was done with fMRIflows (with a temporal low-pass filter at 0.2 Hz; without a low-pass filter looks identical), fMRIPrep, FSL and SPM (from top to bottom) separated for the TR2000 (left), TR1000 (middle) and TR600 (right) dataset. The color value represents the standard deviation value over all subjects. The color scale is the same within a dataset and was set manually to highlight the border effects in gray matter regions. Regions with high inter-subject variability are shown in yellow, while regions with low inter-subject variability are shown in blue. The outline of the brain and subcortical white matter regions is delineated in red and is based on the ICBM 2009c brain template, except for the analysis with SPM where it is based on SPM’s tissue probability map template.

The averaged standard deviation maps after fMRIflows’ and fMRIPrep’s preprocessing are very similar, which is not surprising as fMRIflows uses the same ANTs normalization routine with very similar parameters. The main difference lies in the fact that fMRIflows applies a brain extraction on the functional images as well, which is not performed with fMRIPrep.

#### Temporal signal-to-noise ratio (tSNR) after preprocessing

We computed the voxel-wise temporal SNR according to (Smith et al., 2013) to assess the amount of informative signal contained in the data after preprocessing. This measurement serves as a rough estimate to compare different preprocessing methods but did not allow a direct comparison between datasets, as the tSNR value is a relative measurement that depends highly on the paradigm presented, the initial spatial and temporal resolution of the functional images, as well as the MRI scan sequence specific parameters such as acceleration factors (Smith et al., 2013). Using Nipype’s TSNR routine, we first removed 2^nd^-degree polynomial drifts in each functional image and estimated tSNR maps by computing each voxel’s temporal mean, dividing it by its temporal standard deviation, and multiplying it by the square root of the number of time points recorded in a given run. By averaging the tSNR maps over the population, we get a general tSNR map per preprocessing method for each dataset (see Figure 10).

**Figure 10:**
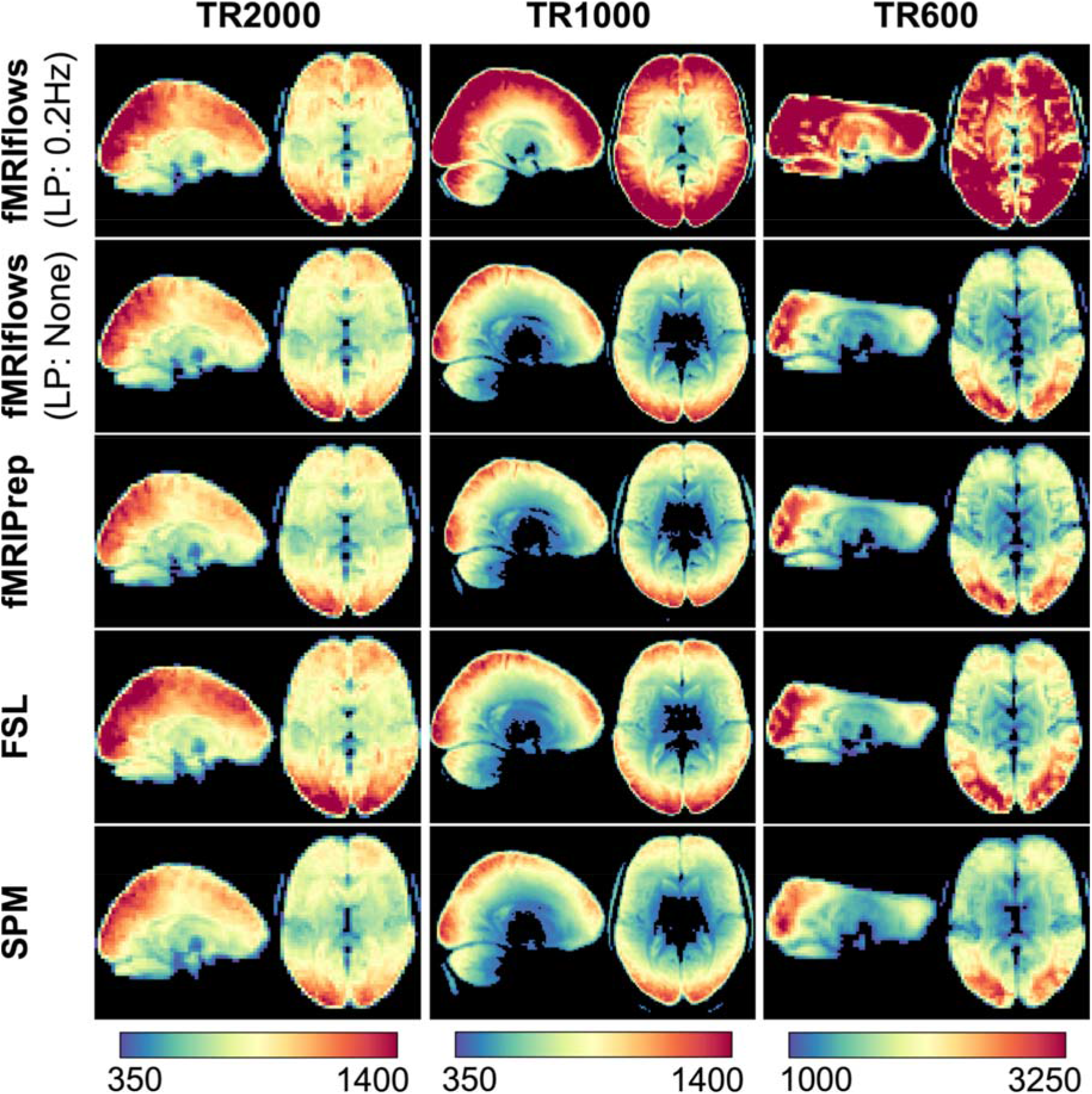
Depiction of temporal signal-to-noise ratio maps of three different datasets, after multiple functional preprocessing approaches. Preprocessing was done with fMRIflows (with and without a temporal low-pass filter at 0.2 Hz), fMRIPrep, FSL and SPM (from top to bottom) separated for the TR2000 (left), TR1000 (middle) and TR600 (right) dataset. The color value represents the tSNR value as computed with the Nipype routine TSNR. The color scale was set manually and differs between datasets, but is held constant between different preprocessing methods.

In general, preprocessing with fMRIflows without a temporal low-pass filter led to similar average tSNR maps as preprocessing with fMRIPrep. Overall, preprocessing with FSL led to slightly increased average tSNR values, while preprocessing with SPM led to slightly decreased average tSNR maps. The additional application of a low-pass filter at 0.2 Hz in all three datasets led to increased tSNR values after preprocessing with fMRIflows. This effect was more pronounced for higher temporal resolution (as in Dataset TR1000 and TR600). The color scales in Figure 10 were set manually so that the fMRIflows (without a low-pass filter) approach shows comparable intensities for the three datasets.

#### Performance check after 1st-level analysis

To investigate the effect of the different preprocessing methods on the 1^st^-level analysis, we carried out a within-subject statistical analysis using Nistats. The activation maps were estimated using a general linear model (GLM). The GLM included a constant term, the stimuli regressors convolved with a double-gamma canonical hemodynamic response function, six motion parameters (three translation and three rotation), and a high-pass filter at 100Hz, represented by a set of cosine functions, and no temporal derivatives. The input data were smoothed using a kernel with a FWHM of 6mm, using a Nilearn routine. The analysis pipelines between the preprocessing methods and datasets were kept as identical as possible and differed only in the number of time points contained in the dataset and the estimated motion parameters. The statistical map for each participant was binarized at z = 3.09, which corresponds to a one-sided test value of *p* < 0.001. The population average of these maps is shown in Figure 11.

**Figure 11:**
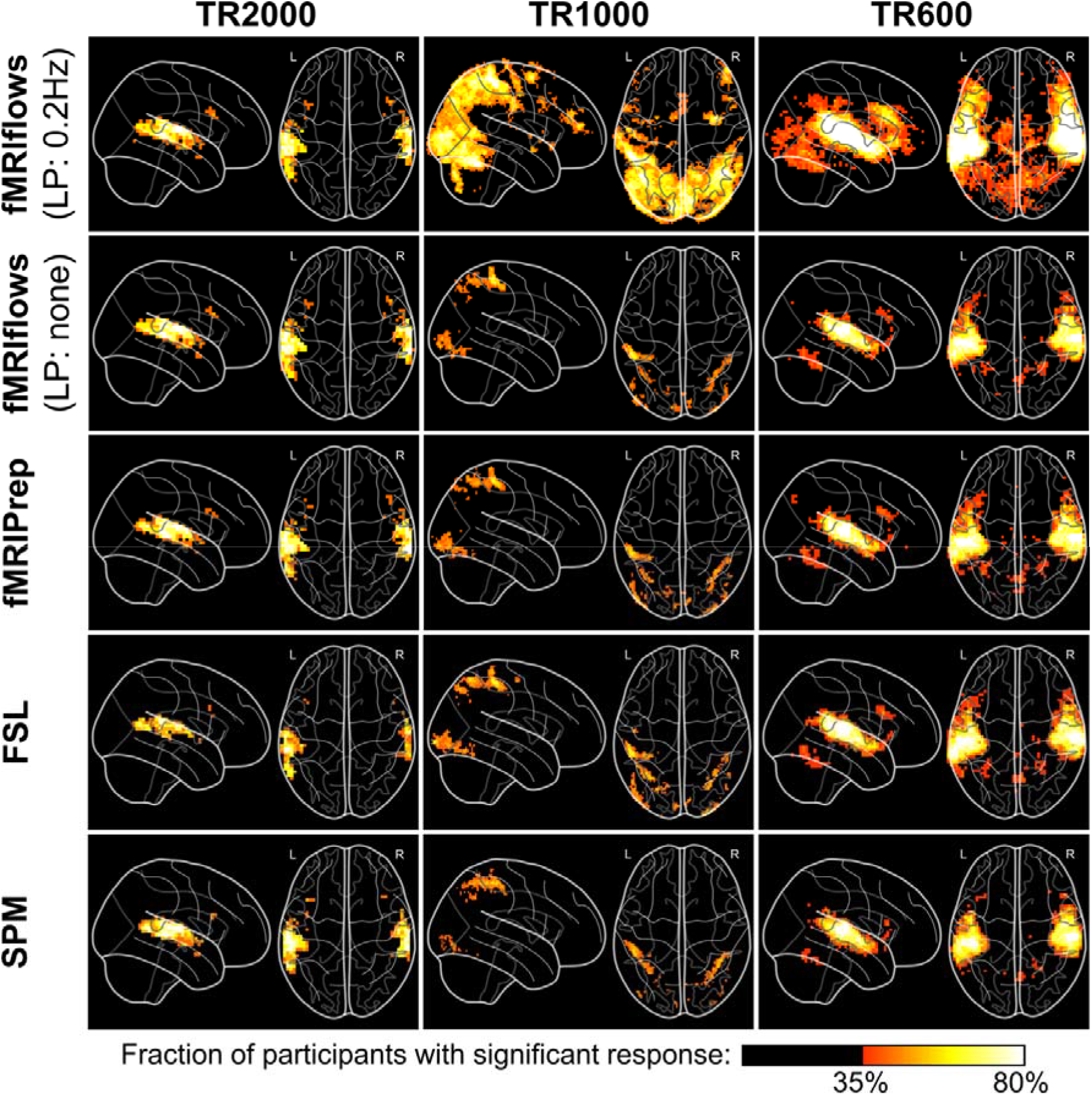
Depiction of binarized 1^st^-level activation count maps, thresholded at *p*<0.001, after multiple functional preprocessing approaches. Preprocessing was done with fMRIflows (with and without a temporal low-pass filter at 0.2 Hz), fMRIPrep, FSL and SPM (from top to bottom) separated for the TR2000 (left), TR1000 (middle) and TR600 (right) dataset. Activation count maps were normalized to the ICBM 2009c brain template. Color code represents the fraction of participants that show significant activation above a *p*-value threshold at 0.001 and corrected for false positive rate (FPR).

The results show that the thresholded activation count maps between the fMRIflows approach without a low-pass filter, fMRIPrep, FSL and SPM do not differ too much between each other, for all three datasets. In contrast to the other preprocessing methods, however, the preprocessing with fMRIflows with a low-pass filter at 0.2 Hz drastically increased the size and fraction value of the thresholded activation count maps, for the datasets TR1000 and TR600. Thus, appropriate temporal filtering increased the statistics for datasets with higher temporal resolution remarkably. For a more detailed comparison between all the toolboxes, see Supplementary Note 7.

#### Performance check after 2nd-level analysis

To investigate the effect of the different preprocessing methods on the 2^nd^-level analysis, we carried out a between-subject statistical analysis using Nistats and computed a one-sample t-test for each preprocessing method and dataset. The unthresholded group-level T-statistic maps of each analysis were then compared to each other on a voxel-by-voxel level using Bland-Altman 2D histograms (Bowring et al., 2018), see Figure 12.

**Figure 12:**
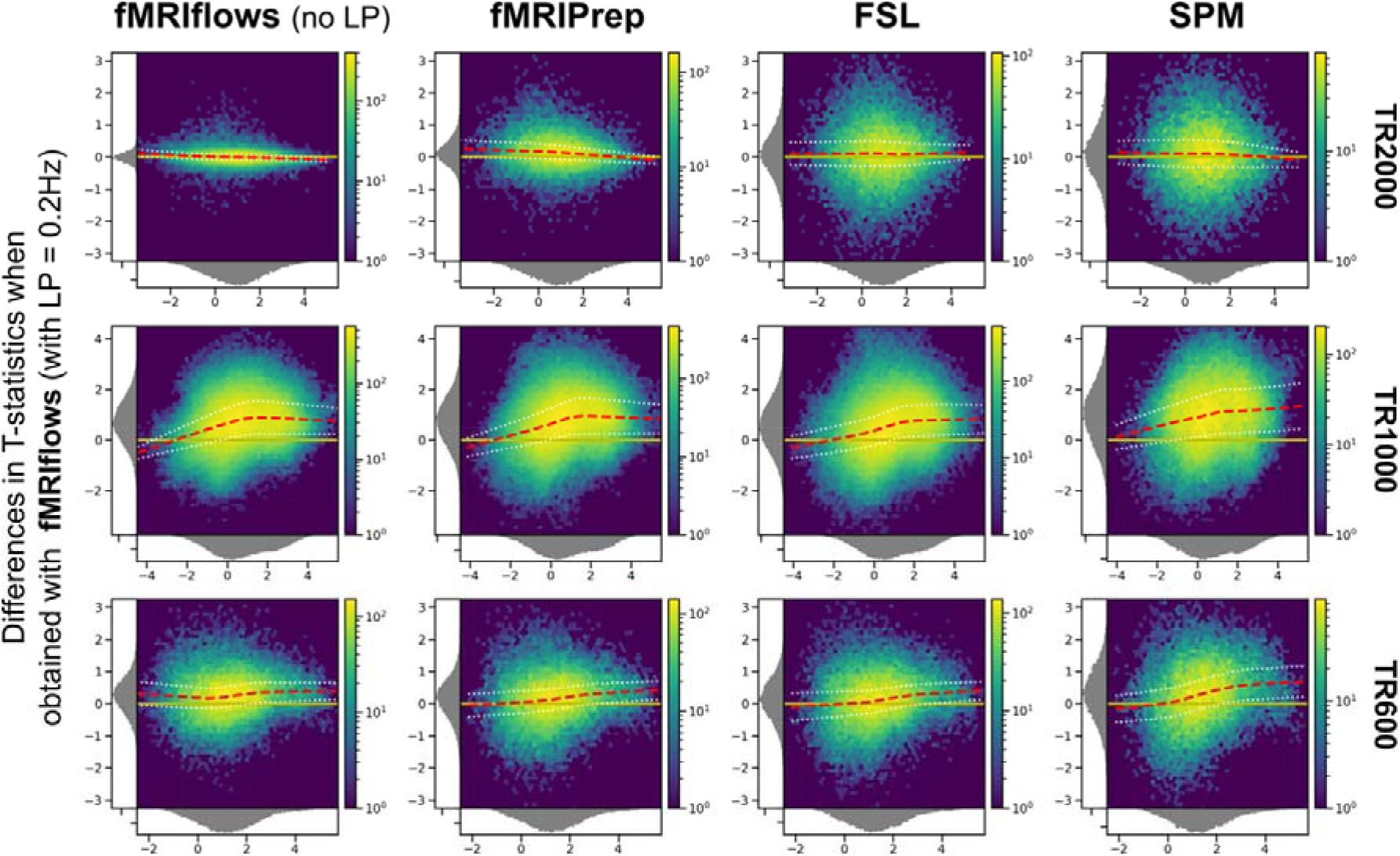
Bland-Altman 2D histograms of three different datasets, comparing unthresholded group-level T-statistic maps between multiple processing approaches. Datasets TR2000 (top), TR1000 (middle) and TR600 (bottom) were used for the comparison. Density plots show the relationship between the average T-statistic value (horizontal) and the difference of T-statistic values (vertical) at corresponding voxels for different pairwise combinations of toolboxes. The difference of T-statistics was always computed in contrast to a preprocessing with fMRIflows using a low-pass filter at 0.2 Hz, while the average T-statistics in horizontal direction investigated the preprocessing with (from left to right) fMRIflows without a low-pass filter, fMRIPrep, FSL and SPM. Distribution plots next to the x- and y-axis depict the occurrence of a given value in this domain. The color code within the figure indicates the number of voxels at this given overlap, from a few (blue) to many (yellow). The yellow horizontal line at zero indicates no value differences between corresponding voxels. The red dashed line depicts the horizontal density average.

The results shown in Figure 12 indicate no pronounced differences between the preprocessing with fMRIflows with a low-pass filter at 0.2 Hz and the other four approaches for the analysis of the TR2000 dataset. Increased variability in the y-direction indicated a decrease in voxel-to-voxel correspondence, which might be explained by different spatial normalization implementations. The fact that the average horizontal density value (red dashed line) is close to the zero line (in yellow) indicated that the different preprocessing methods led to comparable group-level results with the TR2000 dataset. The Bland-Altmann plots for the TR1000 and TR600 datasets showed a clear increase of t-statistic when the preprocessing was done with fMRIflows with a low-pass filter at 0.2 Hz, compared to any other method. This effect was stronger for higher t-values. For a more detailed comparison between all the toolboxes, see Supplementary Note 8.

## 2.3 Discussion

fMRIflows is a fully automatic fMRI analysis pipeline, which can perform state-of-the-art preprocessing, including 1^st^-level and 2^nd^-level univariate analyses as well as multivariate analyses. The goal of such an autonomous approach is to improve the objectiveness of the analyses, maximize transparency, facilitate ease of use, and provide accessible and updated analysis approaches to every researcher, including users outside the field of neuroimaging. While the predefined analysis pipelines help to reduce the number of error-prone manual interventions to a minimum, it also has the advantage of decreasing the number of analytical degrees of freedom available to a user to its minimum (Carp, 2012). This constraint in flexibility is important as it helps to control the variability in data processing and analysis (Botvinik-Nezer et al., 2020). fMRIPrep showed a clear need for such analysis-agnostic approaches and was therefore chosen to provide much of the groundwork for fMRIflows. Our pipeline provides a reliable methodological framework for analyzing functional magnetic resonance imaging (fMRI) data and for obtaining statistical results to be used for advanced multivariate analysis techniques. fMRIflows achieves: 1) high SNR after preprocessing, (2) reproducible within-subject t-statistics, and (3) reproducible between-subject t-statistics. The flexibility for the user to perform both spatial and temporal filtering is particularly important in the context of datasets that had a temporal sampling equal to or below 1000ms or if the statistical output will be used for more advanced analyses, such as MVPA. fMRIflows also improved the overall computation time needed to perform preprocessing and 1^st^ and 2^nd^- level analyses. Indeed, Nipype provides a parallel execution feature of processing pipelines, which is not yet possible with FSL or SPM. fMRIPrep uses the same boost of parallelism but is overall much slower if the default execution of FreeSurfer’s recon-all routine is performed. However, fMRIflows does not yet support parallel computation via a job scheduler on a computation cluster, which is currently possible with fMRIPrep.

In comparison with other neuroimaging software/pipelines like fMRIPrep, FSL and SPM, fMRIflows achieved comparable or improved results in (1) SNR after preprocessing, (2) within-subject t-statistics, and (3) between-subject t-statistics. These results were more obvious in the context of datasets that had a temporal sampling equal to or below 1000ms, and if a low-pass filter at 0.2 Hz was applied.

The inclusion of many informative visual reports allows direct quality control and verification of the performed processing steps, as fMRIflows’ outputs provide a general quality assessment even though it is not as detailed and rigorous as MRIQC (Esteban et al., 2017). In contrast to other software packages, fMRIflows uses an adapted visualization of the carpet plot proposed by (Power, 2017) to highlight the underlying temporal structure and voxel-to-voxel correlations within different brain tissue regions and/or throughout the brain. Such approaches help to observe general signal trends and sudden abrupt signal changes throughout the brain, but the exact implications of these modified carpet plots need to be further investigated.

Being an open-source project, shared via GitHub, facilitates the transparency in the development of fMRIflows. Users can inspect the complete history of the changes and have access to all discussions connected to the software. Code adaptations and additional support for new usage will be proposed by the user community, which will make the adaptation to the newest standards easy and straightforward. In addition to the version-controlled system used on GitHub, a continuous integration scheme with CircleCi will ensure continuous functionality.

Results of fMRIflows’ validation phase 1 suggest that the software is capable of analyzing different types of datasets, independently of the extent of head coverage, original image orientation, and spatial or temporal resolution. By increasing the user base and testing fMRIflows on many more datasets, new adaptations might be required and hidden bugs could emerge. Users can observe any changes done to the software in the future directly on GitHub and are encouraged to state any questions or comments in connection with the software on the community-driven neuroinformatics forum NeuroStars (https://neurostars.org).

Further development of the software will involve (1) moving away from an SPM dependency for the 1^st^ and 2^nd^-level modeling, (2) using the more flexible FitLins toolbox (https://github.com/poldracklab/fitlins) to make the results conform with the BIDS statistical models proposal (BEP002), and (3) implementing an fMRIflows BIDS-App to further improve the toolbox’s accessibility.

## Data Availability

The TR2000 (ds001345, v1.0.1) and TR1000 (ds001734, v.1.0.4) datasets are freely available from OpenNeuro.org. The TR600 dataset will be made available via OpenNeuro.org after publication of the experiments for which it was acquired. Interested parties can in the interim contact the corresponding authors.

## Acknowledgments

We thank the creators of fMRIPrep for providing an excellent starting point and inspiration in the development of fMRIflows. We also thank the developers of PyMVPA, Nilearn, Nibabel and Nistats for bringing the neuroimaging domain into the Python universe, and the developers of Nipype and BIDS for creating a clear framework to execute processing pipelines, as well as the whole neuroimaging open source and science community with its numerous contributors.

## References

Abraham A, Pedregosa F, Eickenberg M, Gervais P, Mueller A, Kossaifi J, Gramfort A, Thirion B, Varoquaux G (2014) Machine learning for neuroimaging with scikit-learn. Front Neuroinform 8:14.

Ashburner J (2009) Preparing fMRI data for statistical analysis. In: fMRI techniques and protocols, pp 151–178. Springer.

Avants BB, Tustison NJ, Song G, Cook PA, Klein A, Gee JC (2011) A reproducible evaluation of ANTs similarity metric performance in brain image registration. Neuroimage 54:2033–2044.

Behzadi Y, Restom K, Liau J, Liu TT (2007) A component based noise correction method (CompCor) for BOLD and perfusion based fMRI. Neuroimage 37:90–101.

Botvinik-Nezer R et al. (2020) Variability in the analysis of a single neuroimaging dataset by many teams. Nature 582:84–88.

Botvinik-Nezer R, Iwanir R, Holzmeister F, Huber J, Johannesson M, Kirchler M, Dreber A, Camerer CF, Poldrack RA, Schonberg T (2019) fMRI data of mixed gambles from the Neuroimaging Analysis Replication and Prediction Study. Scientific Data 6:106.

Bowring A, Maumet C, Nichols T (2018) Exploring the impact of analysis software on task fMRI results. BioRxiv:285585.

Brett M et al. (2018) nibabel: Access a cacophony of neuro-imaging file formats, version 2.3.0.

Caballero-Gaudes C, Reynolds RC (2016) Methods for cleaning the BOLD fMRI signal. Neuroimage:0–1.

Caballero-Gaudes C, Reynolds RC (2017) Methods for cleaning the BOLD fMRI signal. Neuroimage 154:128–149.

Carp J (2012) The secret lives of experiments: methods reporting in the fMRI literature. Neuroimage 63:289–300.

Cox RW, Hyde JS (1997) Software tools for analysis and visualization of fMRI data. NMR Biomed 10:171–178.

Esteban O, Birman D, Schaer M, Koyejo OO, Poldrack RA, Gorgolewski KJ (2017) MRIQC: Advancing the automatic prediction of image quality in MRI from unseen sites. PLoS One 12:e0184661.

Esteban O, Markiewicz CJ, Blair RW, Moodie CA, Isik AI, Erramuzpe A, Kent JD, Goncalves M, DuPre E, Snyder M, Oya H, Ghosh SS, Wright J, Durnez J, Poldrack RA, Gorgolewski KJ (2019) fMRIPrep: a robust preprocessing pipeline for functional MRI. Nat Methods 16:111–116.

Feinberg DA, Moeller S, Smith SM, Auerbach E, Ramanna S, Gunther M, Glasser MF, Miller KL, Ugurbil K, Yacoub E (2010) Multiplexed echo planar imaging for sub-second whole brain FMRI and fast diffusion imaging. PLoS One 5:e15710.

Feinberg DA, Setsompop K (2013) Ultra-fast MRI of the human brain with simultaneous multi-slice imaging. J Magn Reson 229:90–100.

Fischl B (2012) FreeSurfer. Neuroimage 62:774–781.

Fonov V, Evans AC, Botteron K, Almli CR, McKinstry RC, Collins DL (2011) Unbiased average age-appropriate atlases for pediatric studies. Neuroimage 54:313–327.

Friston K, Penny W, Ashburner J, Kiebel S, Nichols T (2006) Statistical Parametric Mapping: The Analysis of Functional Brain Images.

Friston KJ, Williams S, Howard R, Frackowiak RS, Turner R (1996) Movement-related effects in fMRI time-series. Magn Reson Med 35:346–355.

Gorgolewski K, Burns CD, Madison C, Clark D, Halchenko YO, Waskom ML, Ghosh SS (2011) Nipype: a flexible, lightweight and extensible neuroimaging data processing framework in python. Front Neuroinform 5:13.

Gorgolewski K, Esteban O, Schaefer G, Wandell B, Poldrack R (2017) OpenNeuro - a free online platform for sharing and analysis of neuroimaging data. Organization for Human Brain Mapping Vancouver, Canada:1677.

Gorgolewski KJ et al. (2016) The brain imaging data structure, a format for organizing and describing outputs of neuroimaging experiments. Scientific Data 3:160044.

Gorgolewski KJ, Varoquaux G, Rivera G, Schwarz Y, Ghosh SS, Maumet C, Sochat V V., Nichols TE, Poldrack RA, Poline J-B, Yarkoni T, Margulies DS (2015) http://NeuroVault.org: a web-based repository for collecting and sharing unthresholded statistical maps of the human brain. Front Neuroinform 9:8.

Griswold MA, Jakob PM, Heidemann RM, Nittka M, Jellus V, Wang J, Kiefer B, Haase A (2002) Generalized autocalibrating partially parallel acquisitions (GRAPPA). Magn Reson Med 47:1202–1210.

Hallquist MN, Hwang K, Luna B (2006) The nuisance of nuisance regression: spectral misspecification in a common approach to resting-state fMRI preprocessing reintroduces noise and obscures functional connectivity. Neuroimage Nov 15;82:208–25.

Hanke M, Halchenko YO, Sederberg PB, Hanson SJ, Haxby J V., Pollmann S (2009) PyMVPA: A python toolbox for multivariate pattern analysis of fMRI data. Neuroinformatics 7:37–53.

Hunter JD (2007) Matplotlib: A 2D graphics environment. Computing In Science & Engineering 9:90–95.

Jenkinson M, Beckmann CF, Behrens TEJ, Woolrich MW, Smith SM (2012) Fsl. Neuroimage 62:782–790.

Jones E, Oliphant T, Peterson P, others (2001) {SciPy}: Open source scientific tools for {Python}.

Kluyver T, Ragan-kelley B, Pérez F, Granger B, Bussonnier M, Frederic J, Kelley K, Hamrick J, Grout J, Corlay S, Ivanov P, Avila D, Abdalla S, Willing C (2016) Jupyter Notebooks—a publishing format for reproducible computational workflows. Positioning and Power in Academic Publishing: Players, Agents and Agendas:87–90.

Lanczos C (1964) Evaluation of Noisy Data. Journal of the Society for Industrial and Applied Mathematics Series B Numerical Analysis 1:76–85.

Lindquist MA, Geuter S, Wager TD, Caffo BS (2019) Modular preprocessing pipelines can reintroduce artifacts into fMRI data. Hum Brain Mapp:407676.

McKinney W, others (2010) Data structures for statistical computing in python. In: Proceedings of the 9th Python in Science Conference, pp 51–56.

Moeller S, Yacoub E, Olman CA, Auerbach E, Strupp J, Harel N, Uğurbil K (2010) Multiband multislice GE-EPI at 7 tesla, with 16-fold acceleration using partial parallel imaging with application to high spatial and temporal whole-brain fMRI. Magn Reson Med 63:1144–1153.

Notter M, Gale D, Herholz P, Markello R, Notter-Bielser M-L, Whitaker K (2019) AtlasReader: A Python package to generate coordinate tables, region labels, and informative figures from statistical MRI images. Journal of Open Source Software 4:1257.

Oliphant TE (2007) Python for scientific computing. Computing in Science & Engineering 9.

Penny WD, Friston KJ, Ashburner JT, Kiebel SJ, Nichols TE (2011) Statistical parametric mapping: the analysis of functional brain images. Elsevier.

Power JD (2017) A simple but useful way to assess fMRI scan qualities. Neuroimage 154:150–158.

Power JD, Barnes KA, Snyder AZ, Schlaggar BL, Petersen SE (2012) Spurious but systematic correlations in functional connectivity MRI networks arise from subject motion. Neuroimage 59:2142–2154.

Power JD, Plitt M, Kundu P, Bandettini PA, Martin A (2017a) Temporal interpolation alters motion in fMRI scans: Magnitudes and consequences for artifact detection. PLoS One 12:e0182939.

Power JD, Plitt M, Laumann TO, Martin A (2017b) Sources and implications of whole-brain fMRI signals in humans. Neuroimage 146:609–625.

Sengupta A, Pollmann S, Hanke M (2018) Spatial band-pass filtering aids decoding musical genres from auditory cortex 7T fMRI. F1000Res 7:142.

Sladky R, Friston KJ, Tröstl J, Cunnington R, Moser E, Windischberger C (2011) Slice-timing effects and their correction in functional MRI. Neuroimage 58:588–594.

Smith SM et al. (2013) Resting-state fMRI in the Human Connectome Project. Neuroimage 80:144–168.

Smith SM, Jenkinson M, Woolrich MW, Beckmann CF, Behrens TEJ, Johansen-Berg H, Bannister PR, De Luca M, Drobnjak I, Flitney DE, Niazy RK, Saunders J, Vickers J, Zhang Y, De Stefano N, Brady JM, Matthews PM (2004) Advances in functional and structural MR image analysis and implementation as FSL. Neuroimage 23 Suppl 1:S208–19.

Stelzer J, Chen Y, Turner R (2013) Statistical inference and multiple testing correction in classification-based multi-voxel pattern analysis (MVPA): random permutations and cluster size control. Neuroimage 65:69–82.

Stephen Butterworth (1930) On the Theory of Filter Amplifiers. Experimental Wireless and the Wireless Engineer 7:536–541.

Strother SC (2006) Evaluating fMRI preprocessing pipelines. IEEE Eng Med Biol Mag 25:27–41.

Viessmann O, Möller HE, Jezzard P (2018) Dual regression physiological modeling of resting-state EPI power spectra: Effects of healthy aging. Neuroimage 187:68–76.

Yarkoni T et al. (2019) PyBIDS: Python tools for BIDS datasets. Journal of Open Source Software 4:1294.

