## Supplementary material for "fMRIflows: a consortium of fully automatic univariate and multivariate fMRI processing pipelines"

**Supplementary Note 1: Content examples of parameter specification files**

**Content example of file fmriflows_spec_preproc.json**

```
{
    "subject_list_anat": ["01", "02", "03", "04", "05"],
    "session_list_anat": [],
    "T1w_id": "T1w",
    "res_norm": [1.0, 1.0, 1.0],
    "norm_accuracy": "precise",
    "subject_list_func": ["01", "02", "03", "04", "05"],
    "session_list_func": [],
    "task_list": ["multi"],
    "run_list": [1, 2, 3],
    "ref_timepoint": 500,
    "res_func": 2.0,
    "filters_spatial": [["LP", 6.0]],
    "filters_temporal": [[null, 100.0 ], [5.0, 100.0]],
    "n_compcor_confounds": 5,
    "outlier_thresholds": [3.27, 3.27, 3.27, null, null, null],
    "n_independent_components": 10,
    "n_parallel_jobs": 7
}
```

**Content example of file fmriflows_spec_analysis.json**

```
{
    "tasks": {
        "multi": {
            "condition_names": ["cond_01", "cond_02", "cond_03", "cond_04"],
            "contrasts": [
                ["cond_01", [1.0, 0.0, 0.0, 0.0], "T"],
                ["cond_02", [0.0, 1.0, 0.0, 0.0], "T"],
                ["cond_03", [0.0, 0.0, 1.0, 0.0], "T"],
                ["cond_04", [0.0, 0.0, 0.0, 1.0], "T"],
                ["cond_01 > cond_02", [1.0, -1.0, 0.0, 0.0], "T"]
            ]
        }
    },
    "subject_list": ["01", "02", "03", "04", "05"],
    "session_list": [],
    "filters_spatial": [["LP", 6.0]],
    "filters_temporal": [[null, 100.0], [5.0, 100.0]],
    "nuisance_regressors": ["Rotation", "Translation", "FD", "DVARS", "TV"],
    "use_outliers": true,
    "model_serial_correlations": "AR(1)",
    "model_bases": {"hrf": {"derivs": [0, 0]}},
    "estimation_method": {"Classical": 1},
    "normalize": true,
    "norm_res": [1, 1, 1],
    "con_per_run": true,
    "norm_res_multi": [3.0, 3.0, 3.0],
    "analysis_postfix": "",
    "gm_mask_thr": 0.1,
    "height_threshold": 0.001,
    "use_fwe_correction": false,
    "extent_threshold": 5,
    "use_topo_fdr": true,
    "extent_fdr_p_threshold": 0.05,
    "atlasreader_names": "default",
    "atlasreader_prob_thresh": 5,
    "n_parallel_jobs": 7
}
```

**Content example of file fmriflows_spec_multivariate.json**

```
{
    "subject_list": [ "01", "02", "03", "04", "05"],
    "session_list": [],
    "filters_spatial": [["LP", 6.0]],
    "filters_temporal": [[null, 100.0], [5.0, 100.0]],
    "multivariate_postfix": "",
    "clf_names": ["LinearNuSVMC"],
    "sphere_radius": 3,
    "sphere_steps": 3,
    "n_chunks": 6,
    "tasks": {
        "hrf": [
            [["cond_01", "cond_02"],["cond_01", "cond_02"]],
            [["cond_01", "cond_02"],["cond_03", "cond_04"]]
        ]
    },
    "n_perm": 100,
    "n_bootstrap": 100000,
    "block_size": 1000,
    "threshold": 0.001,
    "multicomp_correction": "fdr_bh",
    "fwe_rate": 0.05,
    "atlasreader_names": "default",
    "atlasreader_prob_thresh": 5,
    "n_parallel_jobs": 7
}
```

**Supplementary Note 2: Description of anatomical preprocessing pipeline steps**

**Image reorientation:** To make sure that all images throughout the processing have the same orientation, images are first reoriented with Nipype according to the neurological convention RAS direction convention left-to-right/posterior-to-anterior/inferior-to-superior (RAS+).

**Image cropping:** To make sure that the focus in the anatomical image is on the brain, we use FSL’s robustfov function to remove irrelevant portions of the neck. This is particularly relevant for the later brain extraction step and helps to ensure that the segmentation algorithm focuses on the brain and not on additional body sections.

**Image inhomogeneity correction:** To correct for intensity non-uniformities caused by the inhomogeneity of the bias field during data acquisition, we use ANTs’ N4BiasFieldCorrection algorithm. This step improves the quality of the following image segmentation and is crucial for anatomical images of lower image quality, as they could otherwise fail during the image segmentation.

**Anatomy segmentation:** The image segmentation uses SPM12’s standard image segmentation and provides probability maps for five tissue segments: gray matter (GM), white matter (WM), cerebrospinal fluid (CSF), skull and head. WM masks are partially eroded to make sure that only white matter voxels are used for the mean signal.

**Brain extraction:** The GM, WM and CSF probability maps are combined using Nilearn to create a binary mask which is used to extract the brain. We chose this approach over others as it proved to be more robust and provided the best balance between restriction and inclusion than other algorithms, especially in the context of low image quality.

**Spatial normalization:** As a final step, the extracted brain image is spatially normalized to the ICBM 152 Nonlinear Asymmetrical template version 2009c (Fonov et al., 2011) with ANTs’ antsRegistration algorithm, using nonlinear image registration with a b-spline interpolation.

**Supplementary Note 3: Description of functional preprocessing pipeline steps**

**Image reorientation:** As a first preprocessing step, functional images are reoriented to the neurological convention RAS, using Nipype, to make sure that they have the same orientation as the anatomically preprocessed images.

**Non-steady-state detection:** Afterwards, the first few volumes of each functional run are investigated for non-steady-state volumes using Nipype. If non-steady-state volumes are detected, they are removed before the motion correction is applied.

**Creation of brain masks:** To remove unwanted tissue from functional images a three-step approach was chosen. First, the mean image of the functional run is corrected for intensity inhomogeneity using ANTs’ N4BiasFieldCorrection algorithm. Second, FSL’s BET algorithm is applied to create a binary brain mask. Third, the binary brain mask is dilated by two iterations and holes are filled. This procedure has proven to be optimal in removing non-brain tissue in almost all cases that it was tested on, while ensuring that all brain tissue types are included within the mask. This brain mask is then applied to the functional images before the motion correction step, to make sure that the estimation of the motion parameter is in relation to the brain and not the whole head. The mask is additionally used to restrict the coregistration of the functional images to the anatomical images to the brain tissue and not to the whole head. However, this binary mask is not used to mask the functional images at any time.

**Slice-timing correction:** If specified, slice time correction (Sladky et al., 2011) is performed on the reoriented functional images using SPM, according to the slice onset parameters specified in the BIDS information file and the reference time point mentioned in the fMRIflows JSON specification file. Slice-time correction is applied after the estimation of the head motion as recommended by (Power et al., 2017a).

**Head motion estimation:** Estimation of the motion parameters is performed using FSL’s MCFLIRT algorithm, on the reoriented functional images, after non-steady-states volumes are removed and the brain is masked with the binary mask computed during the previous step. If the user specified a low-pass filter, an additional step is included in the estimation of the motion parameters. This step takes the estimated motion parameters (three rotation and three translation) and applies a Butterworth (Stephen Butterworth, 1930) low-pass filter to each of the six components individually. This step is crucial to guarantee that the motion correction and the temporal filter are orthogonal to each other. Otherwise, previously filtered confounds might be reintroduced at a later step (Hallquist et al., 2013; Lindquist et al., 2019). We are using custom-written Python code to perform this step, using routines from Nilearn, output files from FSL’s MCFLIRT routine and FSL’s avscale routine.

**Intra-subject registration:** The coregistration of the functional image to the anatomical image is based on FSL’s FEAT pipeline and fMRIPrep and uses a two-step co-registration. Both steps use FSL’s FLIRT algorithm. The first step uses the anatomical image to pre-align the mean image from the estimation of the motion parameters, followed by the second coregistration step where the white matter probabilistic image computed with SPM’s segmentation routine is used together with the anatomical image in a boundary-based registration (BBR) approach. The first step uses six degrees of freedom (three rotations and three translations), while the second step uses nine degrees of freedom, adding three scaling degrees. The addition of these scaling degrees was copied from fMRIPrep’s approach (Esteban et al., 2019) and allows the image to be stretched in the direction of the recording. For example, functional images that are recorded in the A-P axis are often squeezed a bit in this direction. Keep in mind that this scaling/stretching is only used for an optimal image coregistration so that the functional images overlay optimally with the white matter boundaries.

**Spatial interpolation:** Once we have the slice-time corrected functional images, computed the correct motion parameters that we want to apply, have the coregistration matrix between the functional and anatomical images, and have the transformation matrix to normalize the anatomical image onto the ICBM 2009c template brain, we can go to the next step and apply all spatial interpolations in a single-shot. The spatial normalization is optional and can also be applied in the later 1^st^-level notebook. But we recommend doing all of those spatial transformations at once to keep the number of spatial interpolations as low as possible. First, we use C3 to combine all the different transformation matrices and apply them to each volume individually with ANTs’ ApplyTransforms routine, using the LanczosWindowedSinc interpolation (Lanczos, 1964). We chose this interpolation over others, as it minimizes the smoothing effect of the interpolation and creates the least artifacts outside the brain volume.

**Creation and application of warped brain masks:** Once the functional images are spatially transformed either into each subject’s structural space using a user-defined voxel resolution, or directly normalized into template space, also using a user-specified voxel resolution, two masks are created to remove irrelevant voxels outside of the brain, such as skull, eyes, and head, as well as voxels that do not have values in most volumes. Those voxels are mostly seen in functional images that only cover part of the brain (slabs) and are introduced through motion at the top or bottom slices of the slab. The first mask is later used to mask the functional image before the application of the temporal filters, while the second mask (confound brain mask) is applied before the extraction of confound signals. For this reason, the first mask should be sensitive enough to keep all voxels within the brain, while the second mask should be specific enough to only keep brain voxels, so that extracted confound curves are only based on nuisance sources within the brain, i.e. the region that we want to clean. The starting point of both masks is again the binary brain mask. For this, we first compute the functional mean image using Nilearn, second correct for intensity non-uniformities using ANTs N4BiasFieldCorrection routine and third apply FSL’s BET routine to create a binary brain mask. This mask is then dilated by two iteration steps, holes are filled up and the mask is then applied to the spatially transformed functional image. The steps to create the first brain mask are as follows: the initial brain mask is first dilated by two voxels, holes are filled up and any voxels that have no signal in more than 1% of volumes are removed. The second brain mask is created as follows: the initial brain mask is first dilated by one voxel, holes are filled, the binary image is again eroded by two iteration steps and afterward, voxels with no signal in more than 5% of volumes are removed.

**Temporal filtering:** The next preprocessing step applies temporal filters if they were specified. The user can decide if they want to apply low-pass, high-pass or band-pass filters, or none. The temporal filtering is performed with AFNI’s 3dBandpass routine. In this step, we also apply the first mask from the previous step to remove voxels that are clearly non-brain tissues. After temporal filtering of the data, the image intensity is normalized in such a way that the white matter distribution peak throughout the functional image is at a value of 10’000.

**Spatial filtering:** The final preprocessing step applies a spatial filter to the functional images if the user specified this. Contrary to other toolboxes, our software allows the application of spatial low-pass, high-pass and band-pass filters. The last one can be especially useful for the preprocessing of data that is later used in a multivariate analysis. (Sengupta et al., 2018) have shown that the correct application of a band-pass filter can drastically improve the prediction accuracy in a multivariate approach. Independently of whether or not a spatial filter was applied, functional images are checked for absolute values above 30’000. If this is the case, images are rescaled to have a maximum absolute value of 30’000. After this, images are stored in integer16 data format to reduce their footprint on the database.

**Supplementary Note 4: Description of 1st-level analysis pipeline steps**

**Data collection:** Preprocessed functional images and model relevant parameters are collected and prepared for the 1st-level analysis.

**Model Specification and estimation:** Model-relevant parameters are combined to create the design matrix needed for the general linear model (GLM) analysis.

**Univariate contrast estimation:** 1st-level contrasts are computed for each subject individually, according to the input parameters specified in the fMRIflows JSON parameter file.

**Optional multivariate contrast estimation:** To create multiple beta contrasts that then can be used for multivariate analysis, fMRIflows computes one beta contrast per condition per run. These beta contrasts are based on the same design matrix like the one for the univariate analysis. This step also creates a CSV-file containing a list of condition identifiers that can be used in the multivariate analysis to label the contrast maps.

**Optional spatial normalization:** The user can spatially normalize the estimated contrasts if they have not been normalized during functional preprocessing. Contrasts need to be normalized, should the user want to use them in a 2nd-level analysis.

**Supplementary Note 5: Description of 2nd-level univariate analysis pipeline steps**

**Model Specification and estimation:** See description in Supplementary Note 4.

**Univariate contrast estimation:** See description in Supplementary Note 4.

**Topological thresholding**: Once the 2nd-level contrast is estimated, fMRIflows uses Nipype’s Thresholding routine to apply first, a voxel-wise threshold, followed by a cluster-wise topological False Discovery Rate (FDR) correction. The user can decide which parameters to use and if topological thresholding should be applied or not.

**Supplementary Note 6: Description of 2nd-level multivariate analysis pipeline steps**

**Data preparation**: To prepare the data for the multivariate pattern analysis (MVPA) with PyMVPA the β-maps from the first level analysis are normalized by voxel-wise z-scoring them. A binary mask is then created to only include voxels in the searchlight analysis that have a value in at least one of the β-maps. This binary mask is then dilated by two voxels and eroded by one to make sure that the mask does not include single voxel holes. The z-scored and masked images are then saved together with a list of corresponding labels in a PyMVPA conform dataset.

**Searchlight classification**: The searchlight classification is performed for each subject individually and is based on the beta contrasts created during the 1^st^-level analysis. The user can specify which binary classification identifiers to use for the training and testing of the classifier. This approach allows the user to look for patterns that distinguish two classes (i.e. when the two classes are the same in training and testing) or to look for patterns that are identifying class differences (i.e. when the two classes are different during training and testing). The second approach allows the investigation of recurrent patterns, even if the stimuli are not the same. For example, training on the differences between visual stimuli of cats and dogs and testing on auditory stimuli of cats and dogs would reveal regions with distinct brain patterns for cats and dogs, independent of the modality of the stimuli. Natively, fMRIflows supports the following classifiers: C- and Nu-SVM classifiers with a linear or radial basis function kernel, Sparse Multinomial Logistic Regression (SMLR) classifier, Gaussian Naive Bayes classifier and k-Nearest-Neighbor classifier. The addition of any other classifiers from PyMVPA or Scikit-Learn is straightforward and can be done very quickly, if needed. Searchlight-specific parameters, such as radius of the sphere and step size can be specified by the user. For each sphere, an *N*-fold leave-one-out cross-validation is performed, where *N* is set by the user, but usually represents the number of runs per subject. The result is one accuracy value per sphere. To account for holes caused by a searchlight step size bigger than one, results between different searchlight spheres are aggregated so that each voxel in the brain mask represents the average accuracy of all searchlight spheres that contained this voxel in their classification. This approach has the additional beneficial effect of increasing the SNR by smoothing the results before the group analysis.

**Group analysis using T-test**: The application of a simple T-test to test the classification accuracies against chance level is not recommended (Stelzer et al., 2013), and users should choose the appropriate group analysis as proposed by Stelzer et al. (2013). Nonetheless, fMRIflows contains this approach, as it can give some preliminary insights into the group results, using extremely reduced computation time.

**Group analysis using random permutations and cluster size control**: The group analysis approach according to Stelzer et al. (2013) includes a permutation step where subject-specific classifications are rerun up to 100 times with randomized labels. In a later step, those randomized control accuracy maps are then combined up to 100’000 times into “false” group averages and help to estimate a voxel-wise null distribution and expected cluster size of the classification accuracy maps, given the dataset and specified classes. The subject-specific searchlight accuracy maps are averaged over all subjects to obtain a group classification accuracy map, which then is tested against the estimated null distribution. The user can additionally specify the strategy for the multiple comparison correction.

**Topological thresholding**: This step is identical to the one used in the 2^nd^-level univariate analysis.

**Supplementary Note 7: Complete 1st-level activation count maps results comparison between toolboxes, separated by dataset.**

**Dataset TR2000:** The comparison between the binarized 1st-level activation count maps computed on the TR2000 dataset shows no clear differences between the toolboxes. The images on the diagonal all seem comparable. The diagonal offset images only show significantly increased activation counts for maps generated with fMRIflows and fMRIPrep, over activation count maps generated with FSL.

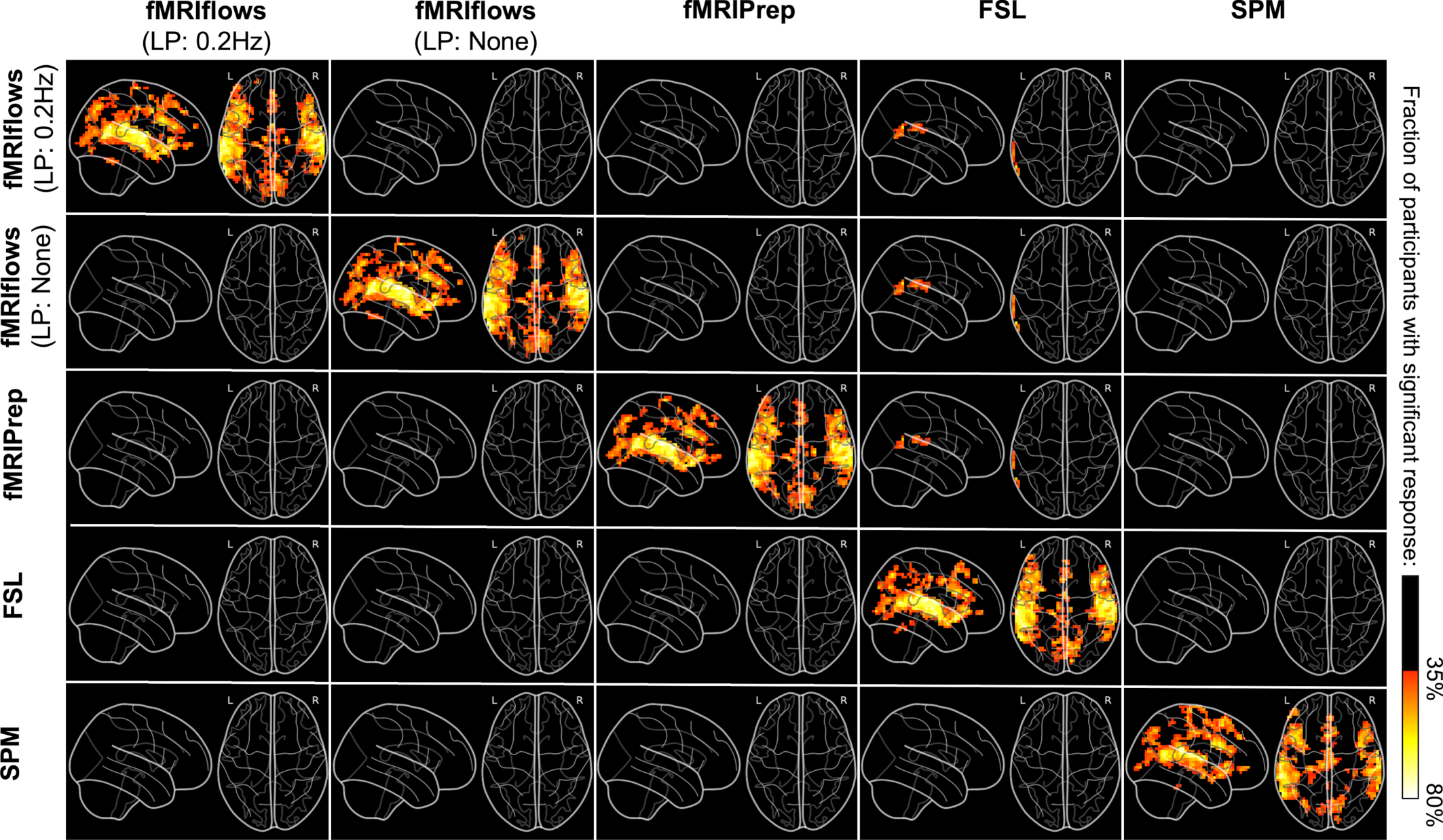

**Supplementary Figure 1**. **Investigation of differences between binarized 1^st^-level activation count maps, thresholded at *p*<0.001, after multiple functional preprocessing approaches analyzing dataset TR2000.** Preprocessing was done with fMRIflows (with and without a temporal low-pass filter at 0.2 Hz), fMRIPrep, FSL and SPM (from top to bottom). The diagonal images represent the original activation count map of the toolbox. The diagonal offset images represent the difference between the horizontal and vertical toolbox. Activation count maps were normalized to the ICBM 2009c brain template. Color code represents the fraction of participants that show significant activation above a p-value threshold at 0.001 and corrected for false positive rate (FPR).

**Dataset TR1000:** The comparison between the binarized 1st-level activation count maps computed on the TR1000 dataset shows clear differences between fMRIflows with a low-pass filter at 0.2 Hz and the other approaches. The images on the diagonal seem overall similar. However, fMRIflows with a low-pass filter at 0.2 Hz shows a clearly increased (and SPM shows a clearly decreased) activation count map. The significantly increased activation count map values between fMRIflows with a low-pass filter at 0.2 Hz and the other approaches, seem to be centered around locations with already increased overlap, indicating that the low-pass filtering improves the overall statistical sensitivity.

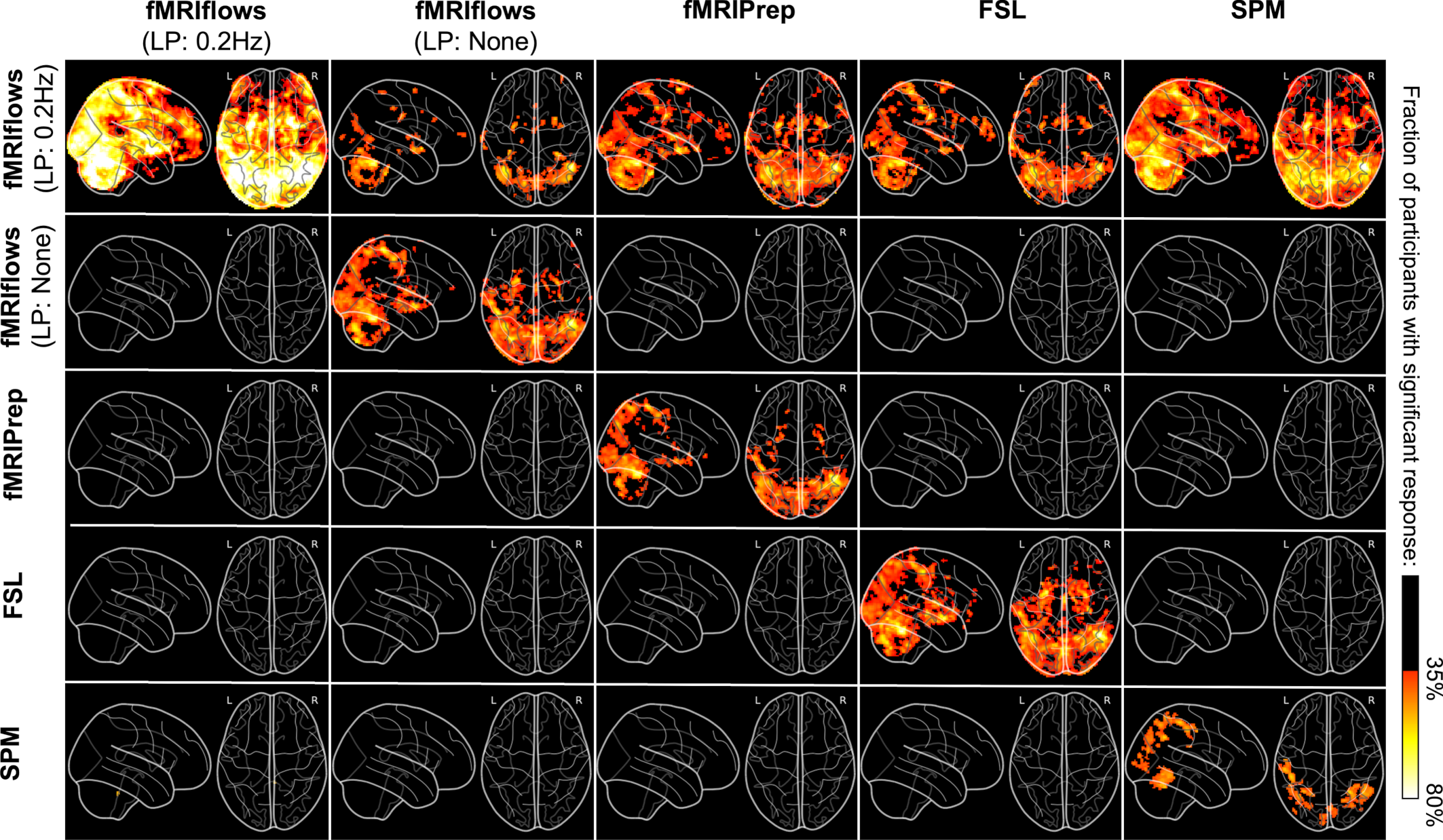

**Supplementary Figure 2: Investigation of differences between binarized 1^st^-level activation count maps, thresholded at *p*<0.001, after multiple functional preprocessing approaches analyzing dataset TR1000.** Preprocessing was done with fMRIflows (with and without a temporal low-pass filter at 0.2 Hz), fMRIPrep, FSL and SPM (from top to bottom). The diagonal images represent the original activation count map of the toolbox. The diagonal offset images represent the difference between the horizontal and vertical toolbox. Activation count maps were normalized to the ICBM 2009c brain template. Color code represents the fraction of participants that show significant activation above a p-value threshold at 0.001 and corrected for false positive rate (FPR).

**Dataset TR600:** The comparison between the binarized 1st-level activation count maps computed on the TR600 dataset shows clear differences between fMRIflows with a low-pass filter at 0.2 Hz and the other approaches. The images on the diagonal seem overall comparable. However, fMRIflows with a low-pass filter at 0.2 Hz shows a clearly increased activation count map. The significantly increased activation count map values between fMRIflows with a low-pass filter at 0.2 Hz and the other approaches, seem to be centered around locations with already increased overlap, indicating that the low-pass filtering improves the overall statistical sensitivity.

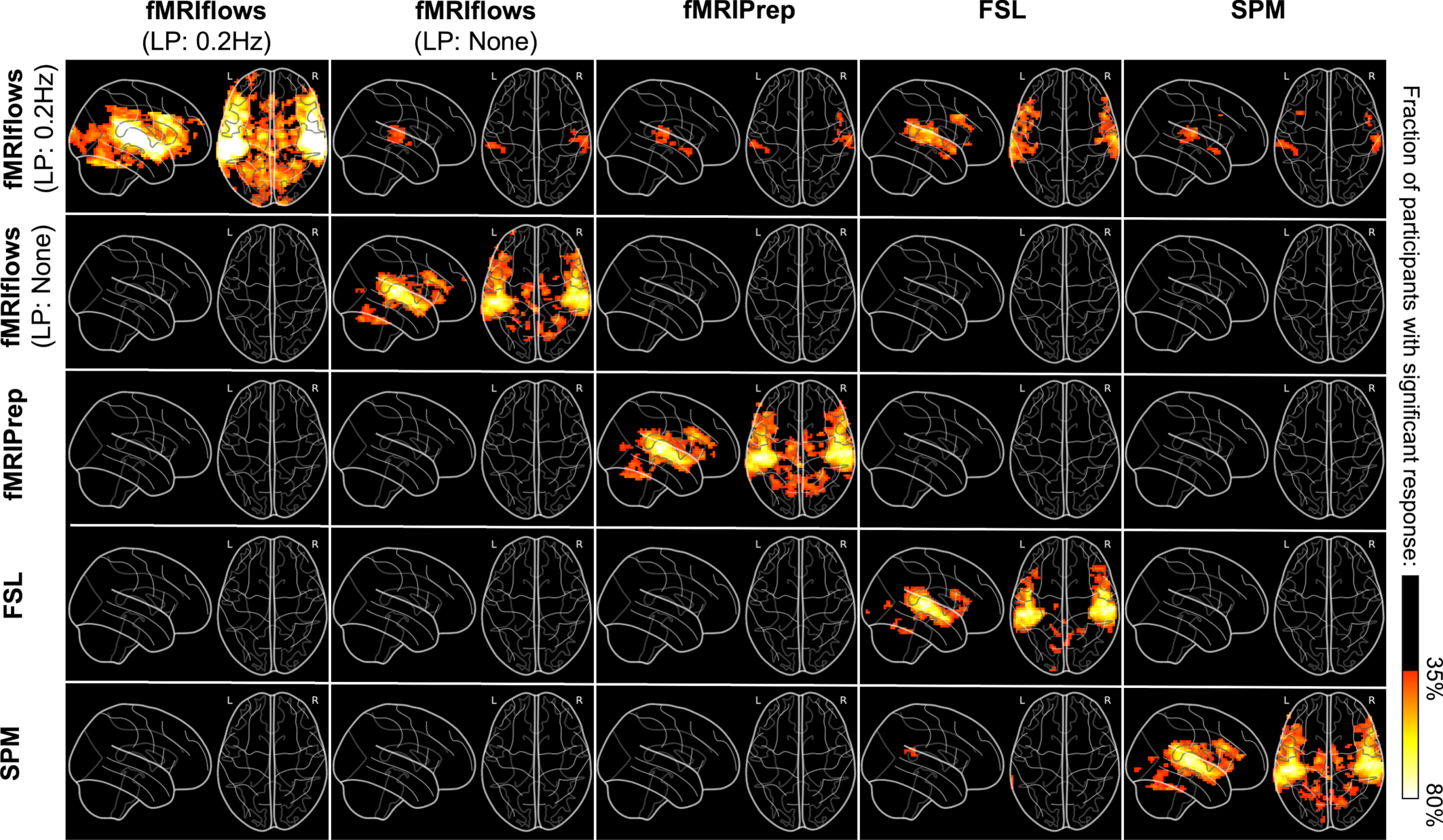

**Supplementary Figure 3: Investigation of differences between binarized 1^st^-level activation count maps, thresholded at *p*<0.001, after multiple functional preprocessing approaches analyzing dataset TR600.** Preprocessing was done with fMRIflows (with and without a temporal low-pass filter at 0.2 Hz), fMRIPrep, FSL and SPM (from top to bottom). The diagonal images represent the original activation count map of the toolbox. The diagonal offset images represent the difference between the horizontal and vertical toolbox. Activation count maps were normalized to the ICBM 2009c brain template. Color code represents the fraction of participants that show significant activation above a p-value threshold at 0.001 and corrected for false positive rate (FPR).

**Supplementary Note 8: Complete group-level T-statistic maps comparison between toolboxes, separated by dataset.**

**Dataset TR2000:** The investigation of group-level T-statistic map differences due to toolbox-specific preprocessing pipelines on the TR2000 dataset does not show any clear differences between the results. The Bland-Altman 2D histograms show no particular trend in the x-direction and are centered around the horizontal zero line.

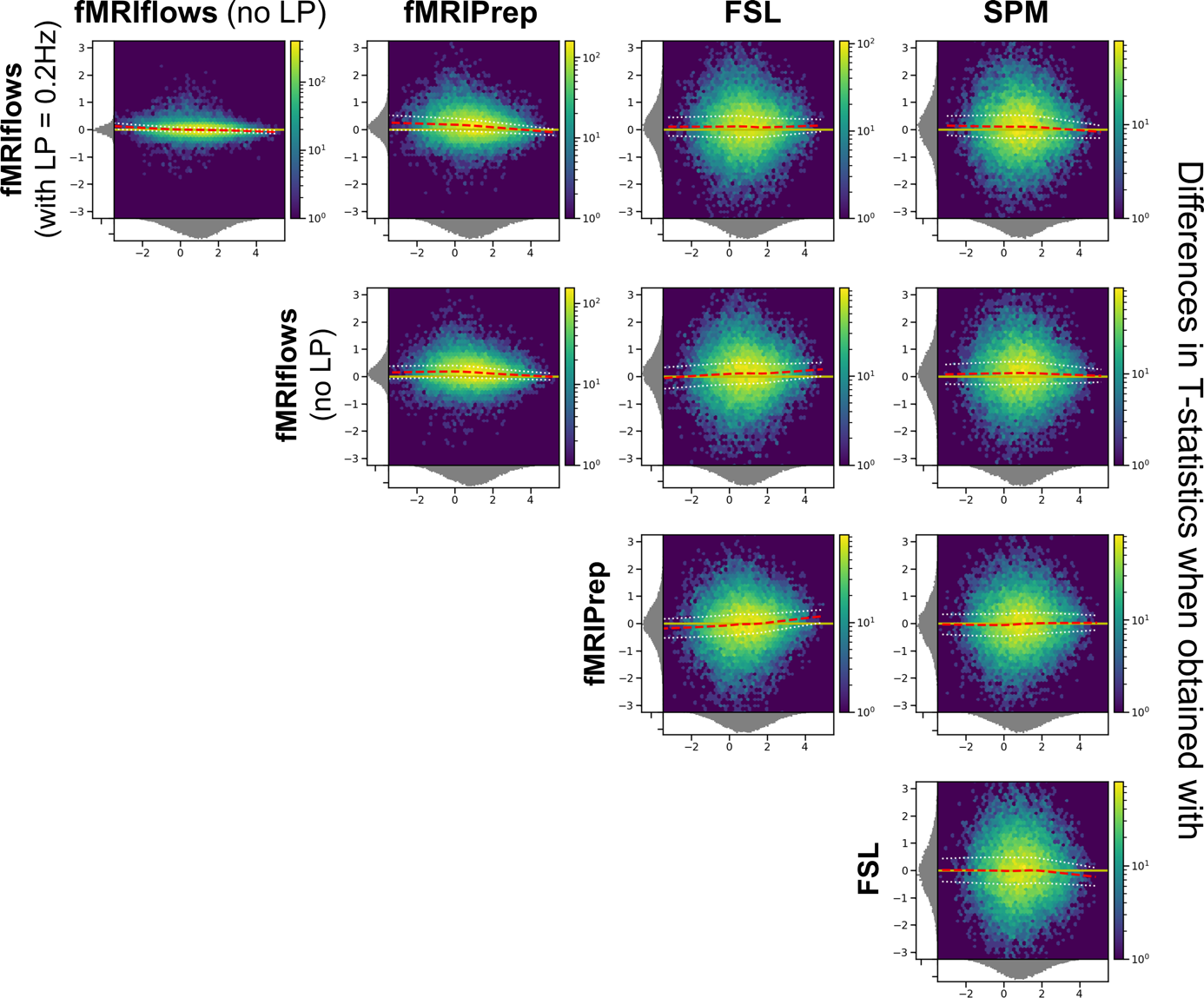

**Supplementary Figure 4: Bland-Altman 2D histograms of datasets TR2000, comparing unthresholded group-level T-statistic maps between multiple processing approaches.** Density plots show the relationship between the average T-statistic value (horizontal) and the difference of T-statistic values (vertical) at corresponding voxels for different pairwise combinations of toolboxes. The difference in T-statistics was computed in contrast to a preprocessing with (from top to bottom) fMRIflows with and without a low-pass filter at 0.2 Hz, fMRIPrep and FSL in respect to a preprocessing with (from left to right) fMRIflows without a low-pass filter at 0.2 Hz, fMRIPrep, FSL and SPM. Distribution plots next to the x- and y- axis depict occurrence of a given value in this domain. The color code within the figure indicates the number of voxels at this given overlap, from a few (blue) to many (yellow). The yellow horizontal line at zero indicates no value differences between corresponding voxels. The red dashed line depicts the horizontal density average.

**Dataset TR1000:** The investigation of group-level T-statistic maps differences due to toolbox-specific preprocessing pipelines on the TR1000 dataset shows clear differences between the results obtained with fMRIflows with a low-pass filter at 0.2 Hz and all the other approaches. The Bland-Altman 2D histograms show clearly increased t-statistic value differences in the first column of the following figure, especially for higher t-values. No clear differences can be seen between the analysis approach fMRIflows without a low-pass filter, fMRIPrep, FSL and SPM.

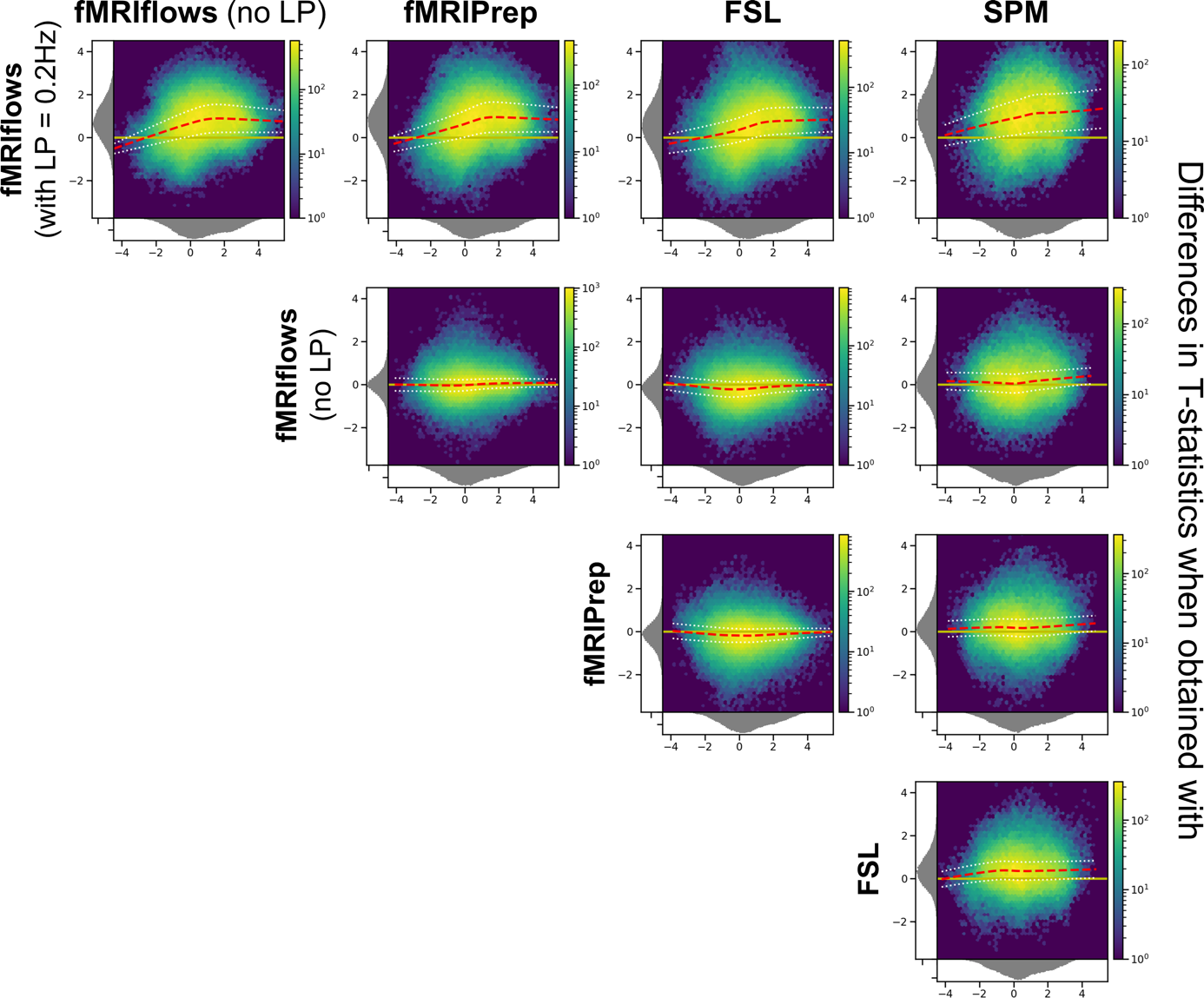

**Supplementary Figure 5: Bland-Altman 2D histograms of datasets TR1000, comparing unthresholded group-level T-statistic maps between multiple processing approaches.** Density plots show the relationship between the average T-statistic value (horizontal) and the difference of T-statistic values (vertical) at corresponding voxels for different pairwise combinations of toolboxes. The difference in T-statistics was computed in contrast to a preprocessing with (from top to bottom) fMRIflows with and without a low-pass filter at 0.2 Hz, fMRIPrep and FSL in respect to a preprocessing with (from left to right) fMRIflows without a low-pass filter at 0.2 Hz, fMRIPrep, FSL and SPM. Distribution plots next to the x- and y- axis depict the occurrence of a given value in this domain. The color code within the figure indicates the number of voxels at this given overlap, from a few (blue) to many (yellow). The yellow horizontal line at zero indicates no value differences between corresponding voxels. The red dashed line depicts the horizontal density average.

**Dataset TR600:** The investigation of group-level T-statistic map differences due to toolbox-specific preprocessing pipelines on the TR600 dataset shows clear differences between the results obtained with fMRIflows with a low-pass filter at 0.2 Hz and all the other approaches. The Bland-Altman 2D histograms show increased t-statistic value differences in the first column of the following figure, especially for high t-values. No clear differences can be seen between the analysis approach fMRIflows without a low-pass filter, fMRIPrep, FSL and SPM.

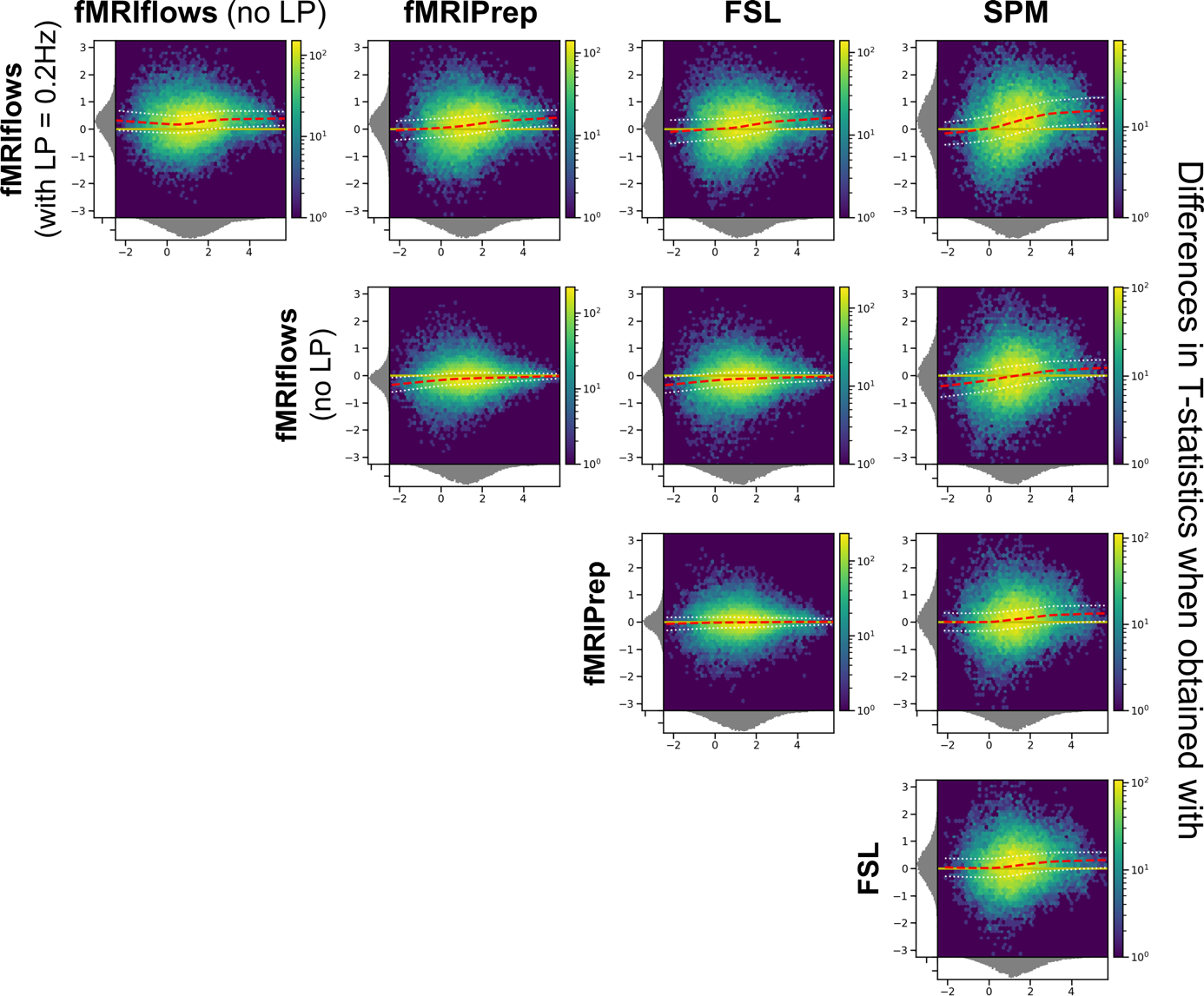

**Supplementary Figure 6: Bland-Altman 2D histograms of datasets TR600, comparing unthresholded group-level T-statistic maps between multiple processing approaches.** Density plots show the relationship between the average T-statistic value (horizontal) and the difference of T-statistic values (vertical) at corresponding voxels for different pairwise combinations of toolboxes. The difference in T-statistics was computed in contrast to a preprocessing with (from top to bottom) fMRIflows with and without a low-pass filter at 0.2 Hz, fMRIPrep and FSL in respect to a preprocessing with (from left to right) fMRIflows without a low-pass filter at 0.2 Hz, fMRIPrep, FSL and SPM. Distribution plots next to the x- and y- axis depict the occurrence of a given value in this domain. The color code within the figure indicates the number of voxels at this given overlap, from a few (blue) to many (yellow). The yellow horizontal line at zero indicates no value differences between corresponding voxels. The red dashed line depicts the horizontal density average.
